# Optical tweezers combined with FRET tension sensor reveal force-dependent vinculin dynamics

**DOI:** 10.1101/2025.11.10.687568

**Authors:** Camille Dubois, Rick I. Cohen, Nada N. Boustany, Nathalie Westbrook

## Abstract

Visualizing and quantifying molecular responses to local forces exerted at cell adhesions is crucial to elucidate how physical forces control cellular behavior. Here, we combine optical tweezers with Förster resonance energy transfer (FRET) microscopy of the vinculin tension sensor, VinTS, to measure the response of vinculin, a key mechanical load-bearing protein, to an applied force. Fibroblasts expressing VinTS formed adhesions on fibronectin-coated, 3μm-diameter, polystyrene beads. As the beads were displaced by the cell, we applied an optical trap to counteract this movement and increase the traction force required by the cell to maintain the bead’s displacement. The median bead displacement after 5 min was ∼200nm in all trapping conditions tested, from zero (no laser) up to 0.26 pN/nm, inducing counteracting forces in the 10-100pN range. To maintain this displacement, vinculin recruitment increased at high stiffness (up to 35% in relative intensity) while vinculin tension increased only moderately in all trapping conditions (1-2% decrease in absolute FRET efficiency). Vinculin recruitment was governed by stiffness rather than the magnitude of the traction force and was correlated with vinculin tension at 0.26 pN/nm but not at lower stiffness. In rare instances, vinculin puncta migrated a few micrometers away from the bead, exceeding the bead’s movement speed while experiencing an increase in both vinculin intensity and tension. Taken together, the results suggest that combining an optical trap with vinculin tension measurements in living cells uncovers novel vinculin dynamics in the presence of a force.

**Why it matters:** Combining optical tweezers with FRET microscopy we measure vinculin tension in conjunction with its recruitment in response to force at adhesions of living cells. We demonstrate that vinculin recruitment is governed by trap stiffness rather than force and that the cell responds to higher stiffness with an increase in vinculin recruitment but less increase in vinculin tension. At high trap stiffness, a correlation between recruitment and tension is observed but not at low stiffness or with no optical trap. Such dynamic measurements, enabled by the techniques presented here, can help elucidate the mechanisms by which cells sense physical forces and the properties of proteins, such as vinculin, which play a fundamental role in cellular behaviors involving tissue growth and repair.

## 1. Introduction

Cell signaling via mechanotransduction is the process by which cells sense and respond to physical forces. Mechanotransduction involving the actin cytoskeleton enables the physiological execution of fundamental dynamic cellular behaviors such as cell movement and cell migration, which are necessary for tissue organization and morphological development (1,2). These processes also play an important role in pathophysiological conditions including wound healing, tissue repair, tumor growth and metastasis (3).

Tension forces within cells, generated by actin polymerization at the leading edge or by actin-myosin contraction, are mediated by the formation of focal adhesions and adherens junctions (4–7) connecting the cell to the extra-cellular matrix (ECM), or to adjacent neighboring cells, respectively. Adhesion complexes initiated when integrins bind to the ECM or when cadherins of neighboring cells bind together, mature under mechanical load into stable adhesions through the activation and recruitment of additional proteins (8).

More than 150 proteins are known to participate in the integrin or cadherin adhesome (9,10). To elucidate the processes by which these proteins are activated and recruited various tools and methodologies have been developed to study the effect of forces on the formation and dynamics of adhesion complexes. Techniques employed to mechanically perturb living cells include varying substrate stiffness (11–19), stretching the substrate (20,21) or applying fluid shear stress to the culture (22). External forces can be applied locally to cells via suction pipets (11,23), optical tweezers (20,24–30) or integrin-coated magnetic beads (31). Disruption of internal cellular forces can also be induced by disrupting actin stress fibers or blocking acto-myosin contractility (11,14,21,23,32–34). Traction force microscopy (16,35,36) and deformable patterned substrates (11,18,37,38) may be used to measure the forces imparted by the cell at focal adhesions while fluorescence microscopy is applied to observe protein recruitment (14,25,39), focal adhesion growth (14,34,40), focal adhesion strength (18,39), adhesion reinforcement (20,41), protein kinase activation (28), YAP nuclear translocation (20), or protein dynamics and actin flow (18,20,42,43), in response to specific mechanical perturbations. Molecular tension probes based on Förster Resonance Energy Transfer (FRET) have added the ability to quantify internal forces on the pN scale on the proteins within focal adhesions, and to do so in a dynamic way (44,45). Mechanical models were also developed to explain protein dynamics at cell adhesions and elucidate cellular behaviors such as migration or sensitivity to the stiffness of the extracellular matrix (12,13,46–48).

In this paper, we combine optical tweezers with fluorescence FRET measurements of the vinculin tension sensor (VinTS). Unlike previous single molecule studies of the vinculin tension sensor involving optical tweezers (44), here we apply the optical tweezers to VinTS inside living cells to probe vinculin tension and its dynamics in response to force. While previous work using optical tweezers has demonstrated adhesion growth and maturation by recruitment of actin and vinculin in response to increasing traction force (24–26,29), dynamic changes in molecular tension in response to an optical trap of varying stiffness have not been previously measured. Tension across vinculin determines the mechanical state of vinculin and leads to the recruitment of additional components to the adhesion site (49). Understanding the dynamic response of vinculin recruitment in conjunction with vinculin tension in response to force is therefore important to elucidate the role of vinculin in strengthening the adhesion complex. To investigate this process, we used human neonatal dermal fibroblasts (HDFn) expressing VinTS and that have initially formed focal adhesions on microbeads coated with fibronectin. As the beads are displaced by the cell, we applied an optical trap to counteract this movement and increase the traction force required by the cell to maintain the bead’s displacement. Over the course of 5 minutes, vinculin is recruited to the adhesions while vinculin tension increases. By varying the optical trap stiffness, our studies show that the level of vinculin recruitment depends on the trap stiffness rather than on the magnitude of the traction force. By contrast, the magnitude of the vinculin tension increase as well as the median bead displacement are not significantly different at the different trap stiffnesses. Thus, adhesion response to higher trap stiffness is mainly governed by higher recruitment of vinculin. Our data also revealed a correlation between vinculin recruitment and vinculin tension at high trap stiffness that is not present at low stiffness. Finally, in rare instances, we observed vinculin puncta moving away from the beads surface. Our results corroborate previous data demonstrating recruitment of vinculin in response to an optical trap but raise new questions regarding the interplay between vinculin recruitment and tension in response to mechanical load.

## 2. Materials and methods

### 2.1. Sample preparation

#### 2.1.1. Beads

Carboxylated polystyrene beads of 3µm or 6µm diameter from Polysciences were functionalized either with fibronectin (FN) (Sigma-Aldrich F0895) or RGD peptide (CellSystems 5020-5MG). For this coating, 10µL of stock beads were incubated with 20µg of poly-lysine in 100µL of PBS for 1h at room temperature, then washed twice with Phosphate-Buffered Saline solution (PBS). The 3µm beads were then incubated with 2µg of FN, and the 6 µm beads with 4µg of FN (or 8µg of RGD), both in 100µL of PBS, for 1h at room temperature. Beads were washed again twice with PBS and resuspended in 100µL of PBS. Fresh beads were prepared a few days before each experiment.

#### 2.1.2. Lentivirus packaging

All procedures were carried out using Biosafety Level 2+ procedures, sterile solutions, and materials unless indicated. The pRRL-VinTS lentivirus vector (50) was generated in HEK293FT cells using protocols with a second-generation plasmid mixture as described previously (51). Details are provided in the Supplementary Methods.

#### 2.1.3. Lentivirus transduction of hTERT-Immortalized Human Foreskin Fibroblasts

Primary normal human neonatal skin fibroblasts (HDFn) from LifeLine Cell Technology (FC-0001) previously immortalized using a pBM14 based vector (52) containing the Open Reading Frame of hTERT (human telomerase reverse transcriptase) (53) were transduced with the 100x concentrated lentivirus containing VinTS. Briefly, subconfluent fibroblast cultures grown in serum medium were rinsed with DMEM/F12 base medium and switched to Optimem containing 10 µg/mL of protamine sulfate. After 1 hour, the lentivirus was added at 1:50, 1:100, and 1:200 dilution. The following day the medium was removed and replaced with a standard serum medium. The expression of VinTS was monitored by microscopy 48 hours post transduction. The 1:50 and 1:100 dilutions resulted in successfully transduced fibroblasts which were subcultured using routine techniques for following tension measurement experiments.

#### 2.1.4. Cell culture

The transduced fibroblasts expressing VinTS were maintained at 37°C, 5% CO2 in high glucose Dulbecco’s Modified Eagle’s Medium supplemented with 10% fetal bovine serum, 1% penicillin/streptomycin, and 1% L-glutamine (all from Gibco). Cells were seeded on 25mm diameter glass coverslips (1.5H Marienfeld Superior), previously cleaned with nitric acid, rinsed 10 times with sterile water, then incubated 1min in methanol, and air dried. Cells were plated at 20,000∕cm2 if imaged the next day or at 10,000-15,000∕cm2 if imaged 2-3 days after plating. Before imaging, cells were mounted in Live Cell Imaging Solution (Invitrogen) and incubated with functionalized beads at 37°C for 45 min, a time long enough for the beads to land on the cells and for focal adhesions to be created around the beads but not too long for internalization to occur. The beads that formed focal adhesions, on which our experiments were performed, were typically at the edge of the cell. The FRET imaging and optical tweezers experiments were conducted at room temperature.

### 2.2. Optical setup

#### 2.2.1. FRET image acquisition

The fluorescence FRET microscopy setup is based on a wide-field sensitized-emission measurement shown in Fig.1 that has been previously described in (54). Briefly, it is composed of two LEDs : a blue LED filtered around 438/24 nm to excite the donor fluorophore mTFP1 and a green LED filtered around 520/15 nm to excite the acceptor mVenus. The illumination intensity of each LED in the sample plane can be adjusted independently. They are chosen based on the half-life of the fluorophores which are set to 150 s for mTFP1 (33.0 mW∕mm2 for LED D) and 40 s for mVenus (11.6 mW∕mm2 for LED A). Half-life of the two fluorophores as a function of illumination intensity is shown in Fig. S1. Fluorescence is collected through a microscope objective (Nikon Plan Fluor 100x, NA = 1.3, oil immersion) and the donor and acceptor emissions are detected simultaneously on two sCMOS cameras (Basler acA2040-55 μm, 2048 x 1536 pixels, 3.45 μm pixels, quantum efficiency 65% at 550 nm, cameras 1 and 2 in Fig. 1), one for the donor and one for the acceptor. An additional camera (camera 3 in Fig. 1) provides a phase contrast image of the sample. An acquisition is automated without any mechanical movement. A FRET acquisition sequence consists of 2 steps: donor is excited with LED D and two images are acquired simultaneously, IDD (Donor excitation and Donor detection) on camera 1 and IDA (Donor excitation and Acceptor detection) on camera 2. Then, the acceptor is directly excited with LED A and the IAA image (Acceptor excitation and Acceptor detection) is acquired on camera 2. All the exposure times are 2s.

**Figure 1:**
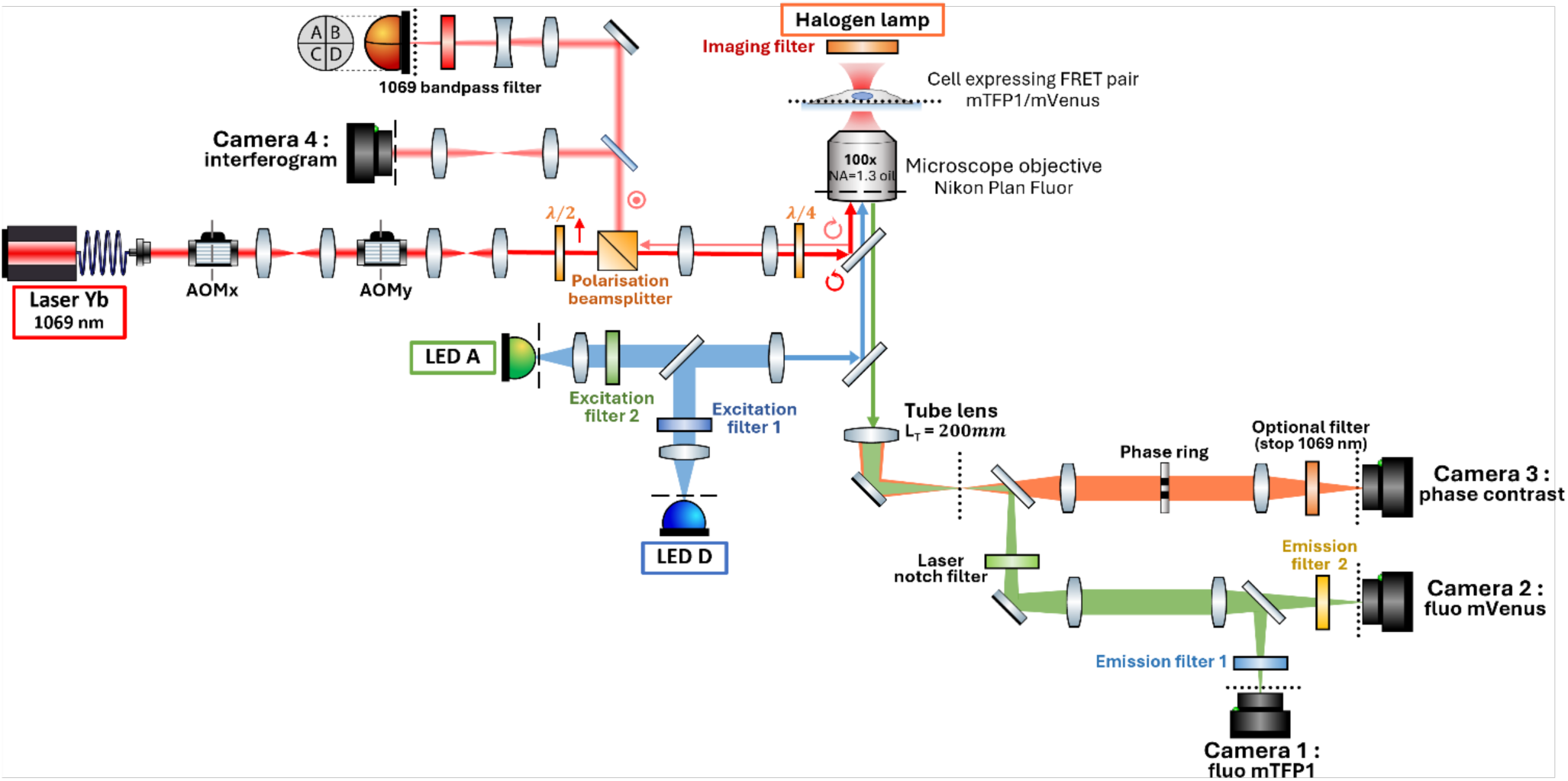
Experimental setup. Conjugate planes to the sample are indicated by dotted lines, those conjugate to the back focal plane of the objective by dashed lines. Filters used for FRET are 438/24 for Excitation 1, 520/15 for Excitation 2, 482/35 for Emission 1, 562/40 for Emission 2. For more details on filters and dichroic mirrors, see (54).

#### 2.2.2. FRET efficiency calculation

The three raw images are processed with a Matlab (The MathWorks Inc.) routine. First, each image is background corrected and flat-field corrected. Then, a registration procedure based on a transformation matrix allowing for translation, rotation and scaling is applied to the IDD image so that it matches pixel-by-pixel with the images taken on the other camera. Fluorescence signals coming from the cells are isolated from the background by thresholding. Fluorescence signals isolated and corrected from each image are called *DD*, *DA* and *AA*.

Bleedthrough between channels was previously measured with cells transfected with acceptor only or donor only: the direct excitation of the acceptor when illuminating with the blue LED at 440 nm gave a ratio a = 0.169 between *DA* and *AA* and the donor fluorescence that passes through the acceptor detection channel gave a ratio d = 0.413 between *DA* and *DD*. These lead to a corrected FRET intensity: *Fc* = *DA* − *aAA* − *dDD*.

Using the calibration procedure described in (54), a G factor has been determined using cells expressing a standard FRET construct, TSMod, whose FRET efficiency has been measured independently by fluorescence lifetime microscopy (FLIM). Therefore, a quantitative FRET efficiency is given by:

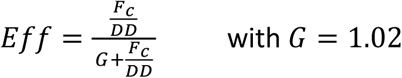

#### 2.2.3. Optical tweezer setup

The optical tweezer is based on an Ytterbium fiber laser at 1069 nm (Keopsys, TEM00, linearly polarized) delivering up to 5 W at the output of the fiber. The beam is expanded by two afocal telescopes to cover the entrance pupil of the objective. The laser waist measured at the focal point is *ω*0,*x* = 675nm and *ω*0,*y* = 470nm.

Two orthogonal acousto-optic deflectors (AOD) (Intra Action Corp. DTD 274HA6) control the displacement of the trap in the transverse plane. The AODs are conjugate to the back-focal plane of the objective to get a pure translation of the trap in the sample plane.

The retroreflected light is separated from the incident beam by polarization and is imaged on a quadrant photodiode (QPD, SPOT-9DMI, OSI Optoelectronics). This detection samples the bead position at 62.5 kHz. Camera 4 in Fig. 1 is conjugate to the back-focal plane of the objective and images the interference pattern between the reflections at the bead bottom surface and at the top surface of the coverslip. Centering the rings in this interferogram is used to center precisely a bead at the center of the trap. The diameter of these rings also reports on the height of the bead with respect to the coverslip (55).

#### 2.2.4. Optical tweezer calibration

##### 2.2.4.1. Lateral position of the bead

Output signals of the QPD are normalized dimensionless signals and are converted into a metric position measurement using a conversion factor C. This factor is obtained using the step-response method : the trap is moved rapidly by a known distance in x and y and the QPD signal is measured as the bead comes back into the trap. The relative voltage of the QPD as a function of the step amplitude is linear over 400 nm around the trap center with a calibration factor *C* = 1.0 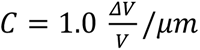/*μm* in both directions x and y (see supplementary Fig. S2).

##### 2.2.4.2. Trap stiffness

The trap stiffness *k* is estimated from the motion of a bead trapped in water using the power spectrum analysis and the step-response method (55). The two methods are used in parallel on the same beads to ensure a reliable calibration. Stiffnesses obtained with the two methods are in close agreement and increase linearly with laser power with a slope of 0.71 pN/nm/W for the 3µm diameter beads used in the trapping experiments (see supplementary Fig. S3). We adjusted the trap stiffness using laser power, with a limit at 370 mW (at the entrance of the objective) to prevent sample damage. The trap stiffness was checked to be linear over the 400nm range of linearity of the QPD. Two power levels were used, 370mW or 180mW, corresponding to trap stiffnesses of 0.26 pN/nm and 0.13 pN/nm respectively, for 3 µm beads.

#### 2.2.5. Trapping and image acquisition sequence

Fig. 2 shows the time sequence of our experiments. Once a bead with puncta of vinculin is identified, it is centered on the laser spot at low power, i.e. less than 10mW (Fig. 2a). The laser is rapidly turned to its nominal power, within less than 10 seconds (Fig. 2b). As the actin reorganization within the cell displaces the bead along its membrane, the laser will induce a counteracting force on the bead (Fig. 2c). From the displacement value x of the bead from its initial position, we infer the counteracting force of the optical trap on the bead as *F*(*t*) = *kx*(*t*), where the stiffness k of the trap can be varied with laser power.

#### 2.2.6. Image processing

The focal adhesions on the beads are extracted by manually drawing an annular region around the bead that includes the focal adhesions. This region of interest is adjusted for every acquisition: the annulus has a central radius between 40 and 55 pixels (corresponding to 1 to 1.4 μm) and a width between 20 and 35 pixels (i.e. 0.5 to 0.9 μm). The center of the annulus is shifted linearly over the time sequence to follow the bead. For each bead, focal adhesions are delimited angularly within this annulus. The number of focal adhesions per bead varies from 1 to 3, and their angular size ranges from 30 to 150° over all the focal adhesions analyzed. AA intensity and FRET efficiency values are averaged over each focal adhesion. Focal adhesions with initial AA intensity below 20 were excluded from the analyses as the signal-to-noise ratio below this intensity level results in a large uncertainty in the calculated FRET efficiency. To quantify the changes over time, the first image taken with the laser on (time t= 1min) is our reference in the analyses for the intensity and FRET efficiency measurements. The AA intensity values are normalized to the AA intensity at t= 1min, and FRET efficiency is reported over time as the difference between the FRET efficiency at time t and the FRET efficiency at t= 1min. Statistical tests were conducted in GraphPad Prism (version 10.0.0 for Windows, GraphPad Software, Boston, Massachusetts USA.)

**Figure 2:**
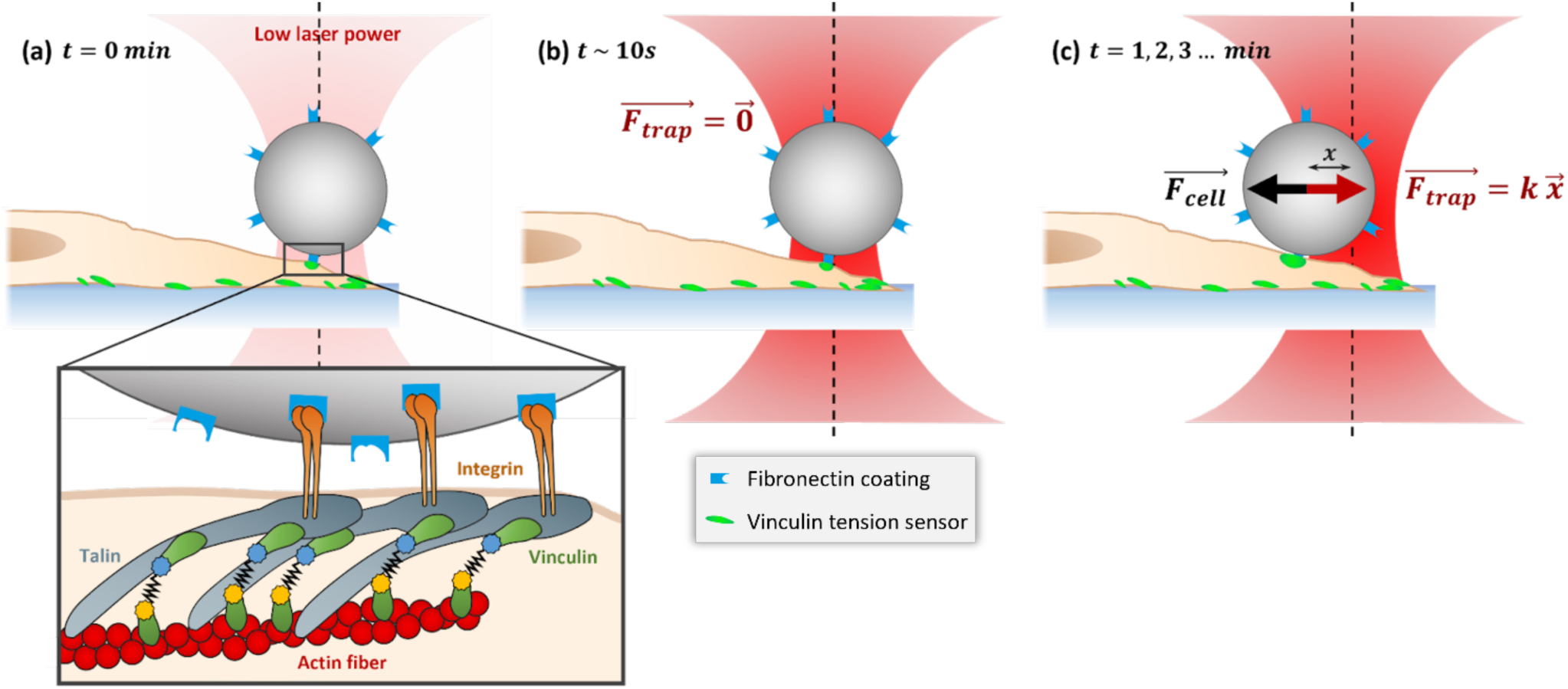
Principle of the application of the optical trap force on the cell. (a) A bead showing puncta of vinculin fluorescence is initially centered on the laser focus at low power using the interference pattern on Camera 4 (see Fig. 1). The first FRET sequence (consisting 3 images DA, DD and AA) is acquired at t=0 and is the first acquisition of the time-lapse. The inset shows the focal adhesion complex with the main proteins involved, and a FRET tension sensor inserted within vinculin, whose head and tail are bound to talin and actin, respectively. (b) The laser power is increased to the desired level immediately after the end of the first FRET acquisition. At that point, the bead is still centered so it does not experience a force from the trap. (c) As the cell displaces the bead, a force is exerted by the laser on the bead equal to *F* = *k x* with k the trap stiffness and x the distance of the bead from its initial position. FRET sequences are recorded every minute.

### 2.3. Confocal imaging

Confocal microscopy was used to visualize the focal adhesions with additional actin fluorescent labelling. Actin in fibroblasts expressing VinTS were labeled using CellLight Actin-RFP, BacMam 2.0 (Invitrogen C10583) following the manufacturer’s protocol. On the next day, cells were incubated with 6µm fibronectin coated beads for 45 min and then imaged. The mVenus within VinTS was excited with a laser at 514 nm and imaged in a bandwidth of 516-597nm. The RFP on actin was excited with a laser at 594nm and imaged in a bandwidth of 596 to 695 nm. Confocal microscopy was performed on a commercial laser scanning microscope (Zeiss 780 LSM) at the Rutgers BME High Resolution Microscopy Core Facility. 3D image rendering was performed using ImageJ Fiji software (56).

## 3. Results

### 3.1. Focal adhesion formation on beads

Fig.3 displays confocal images of a fibroblast cell, highlighting distinct puncta of vinculin around fibronectin-coated beads. These puncta are separate from the focal adhesions found at the surface of the coverslip. This separation is illustrated in the xz and yz profiles presented alongside the right panel in Fig. 3a. The diffuse distribution of vinculin within the cell effectively delineates the cell membrane, confirming that the membrane surrounds the bead, as evidenced by the xz and yz profiles and the 3D renderings in Fig. 3b. Actin stress fibers are clearly visible on the coverslip surface. Additionally, some actin filaments also emerge from focal adhesion points at the level of the bead. Focal adhesions linked to actin filaments were observed on multiple 6 µm beads (more examples in supplementary Fig. S4(a)) suggesting that the puncta we observe and to which vinculin is actively recruited, involve interaction with fibronectin. Results obtained with beads coated with RGD peptide are shown in supplementary Fig. S.4(b) for comparison. In the case of RGD beads, we did not observe the same vinculin puncta or actin organization around the bead showing that the formation of the observed vinculin puncta requires fibronectin.

**Figure 3:**
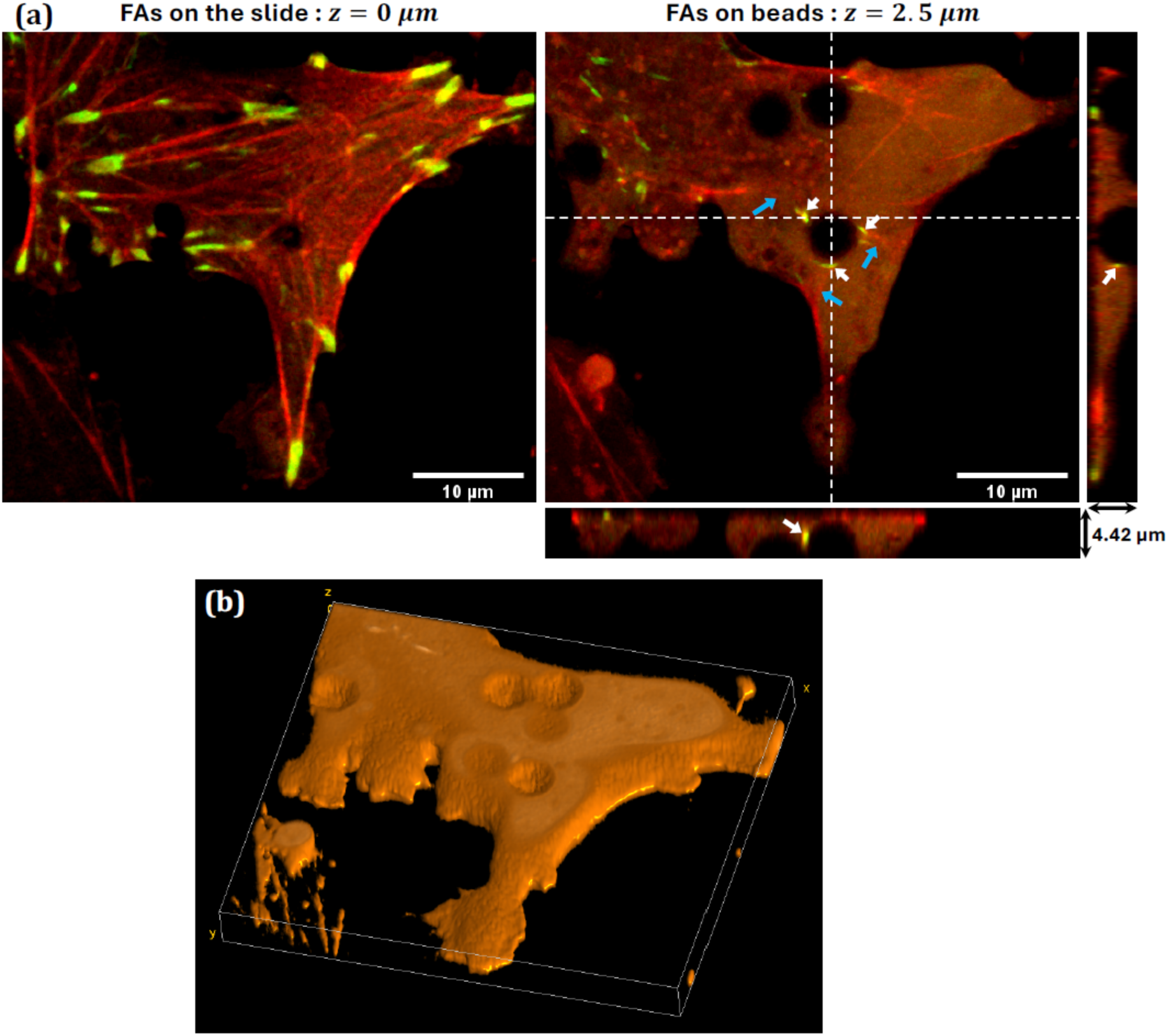
Confocal images of fibroblast cells after 45 min of incubation with 6µm beads coated with fibronectin. (a) Vinculin is marked using the tension sensor VinTS in green and actin is in red. Focal adhesions (FAs) on the coverslip are visible at *z* = 0 *μm* (left). At the height of the beads, *z* = 2.5 *μm* (right), three vinculin puncta (marked by white arrows) are present on the surface of the bead. Actin filaments connected to these FAs are marked by blue arrows. Horizontal and vertical cross-sections along the white dashed lines, corresponding to the xz and yz profiles displayed below (xz) and on the right (yz), show that the vinculin puncta are present at a specific height (z). (b) Volumetric reconstruction of the cell based on VinTS fluorescence, expressed over the whole cytoplasm. The spherical depressions correspond to the positions of the beads. The beads appear slightly embedded in the membrane.

### 3.2. Time response of vinculin recruitment and tension in response to the optical trap

#### 3.2.1. Measuring intensity and FRET efficiency as a function of time on one bead

A typical example of the data recorded for one bead is shown in Fig. 4. For higher stiffness in optical trapping (see stiffnesses for different bead diameters in supplementary Fig.S3) we used 3µm beads, which exhibit similar vinculin puncta as the 6µm beads. The phase contrast image in Fig. 4a shows that several beads are located on the cell, but one clearly presents accumulation of vinculin in the fluorescence image (Fig. 4b), corresponding to initial adhesions. We position this bead at the center of the trapping laser at low power. Fig. 4c shows the intensity (sum of the 3 images DD, DA and AA) and Fig. 4d the FRET efficiency around the bead as a function of time. On the intensity images, we see that the two focal adhesion areas, on the right and left sides of the bead, both increase in intensity. Other examples of intensity images, recorded on different beads and cells, are shown in supplementary Fig. S5. Given the similar evolution of all focal adhesions on the same bead and the fact that they are subjected to the same optical trapping force, we averaged the values over all the focal adhesions for a given bead (between 1 and 3 FAs per bead).

**Figure 4:**
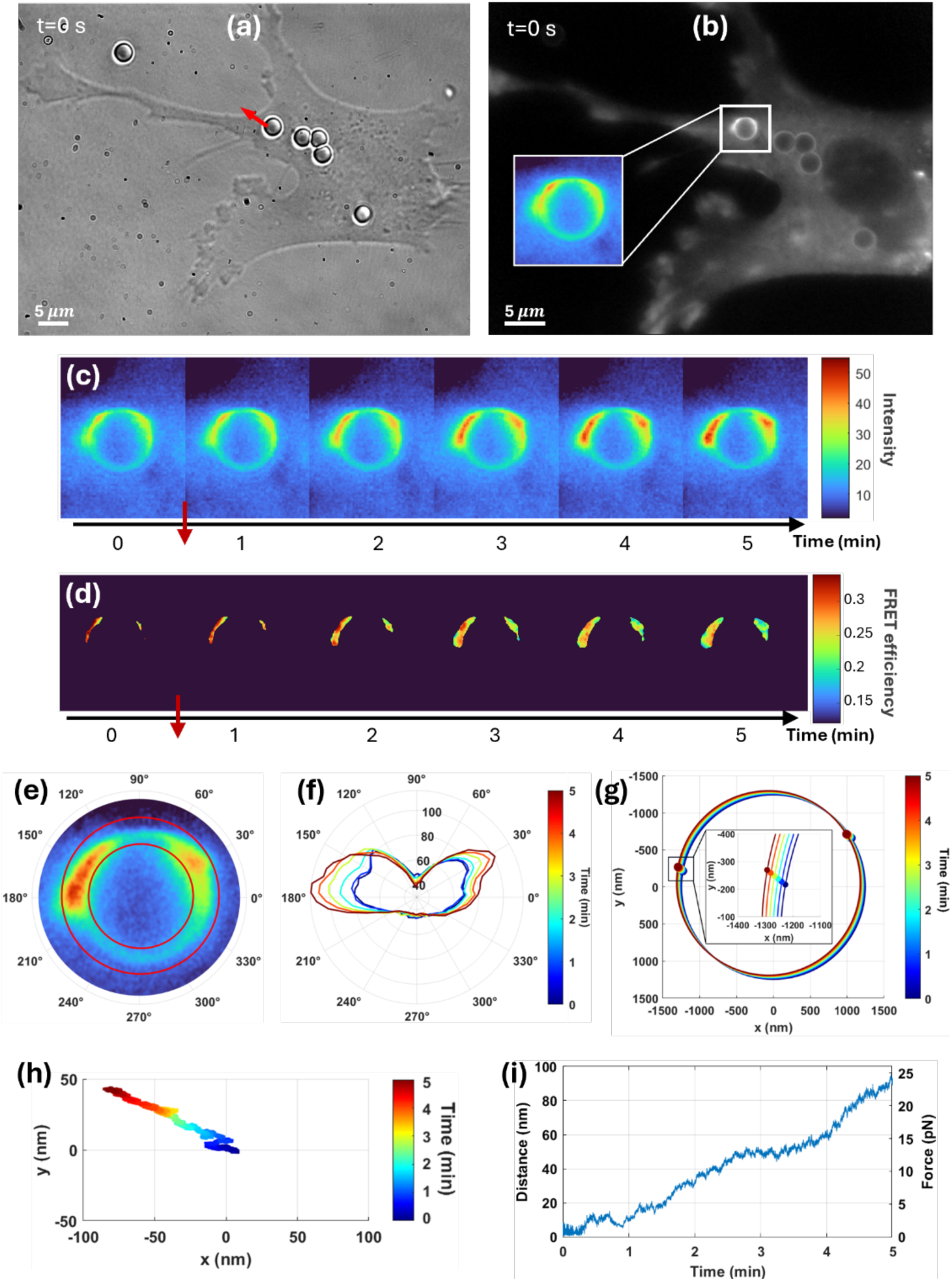
Signals recorded for one bead, shown as an example. Figures (a) and (b) are respectively the phase contrast and fluorescence images of a cell incubated with 3µm FN beads before the beginning of the laser trapping. This direction of motion of the bead is shown as a red arrow in panel (a). (c) Fluorescence images of the focal adhesions around the trapped bead recorded every minute after the laser is turned up to nominal power. The 3 images DD, DA and AA have been summed to increase contrast. AA intensity represents typically ⅓ of the total intensity. (d) Corresponding FRET efficiency images of the adhesions around the trapped bead. (e) Zoom on the fluorescence image at t=5 min, showing the annular region defined for the quantitative analysis of AA intensity and FRET efficiency. Two focal adhesions have been delimited on this bead, one at 30°±15° and 170°±20° (f) Polar plot of the AA intensity averaged over the width of the annular region shown in panel (e). Color codes for time. (g) Displacement of the annular region over the 5 recorded images: the circle segments represent the bead’s position, and the dots the center of the focal adhesion, both as a function of time. (h) Trajectory of the bead recorded on the quadrant photodiode and (i) displacement and force measurement calculated from this trajectory using the calibration described in the Methods. The direction and displacement of the bead in panel (g) are consistent with those measured from the trajectory in panel (h).

As explained in the Methods (Section 2.2.6), for the quantitative analysis, we draw an annulus on the fluorescence image of the bead that encompasses the focal adhesion regions. Fig. 4e shows this annular region for this particular bead at time t = 5 min, and Fig 4f shows the corresponding polar plot of the AA intensity as a function of time. Once we have manually defined the annulus around the bead on the two images at t=0 and t=5 min, we verified that the bead displacement was linear for images at intermediate times (see Fig 4g with the spatial location of the annulus on all 5 images in the xy plane, showing both amplitude and direction of motion). This measurement of the displacement has been used in all our subsequent analyses. Its precision is limited by the camera pixel size (26nm in the sample plane). The displacement deduced from the QPD signal, as shown in Fig 4h, agrees with the amplitude and direction of motion extracted from the fluorescence images. Even though the QPD signal should give us a continuous and more precise measurement of the displacement, transient parasitic fringes in the reflected laser spot recorded on the QPD made it more reliable to use the displacement value extracted from the fluorescence images.

#### 3.2.2. Initial conditions

To assess the initial state of the focal adhesions observed on the different beads, we analyzed the initial intensity and FRET efficiency before applying the laser force. A scatter plot of the initial FRET efficiency values (Fig. 5) plotted against the initial mVenus intensity (AA channel), shows that initially, vinculin is under different levels of tension on the different beads. FRET efficiency values span all the tension sensor’s range: from high efficiency around 30% where vinculin is not under tension down to 15% where vinculin is subjected to high stretch, with most points under 28%, i.e. under some tension. The initial FRET efficiency is not correlated (R2=0.027, p=0.29) with the initial AA intensity (colors indicate the laser trap stiffness that will later on be applied on each bead) consistent with adhesions in different initial states of assembly.

#### 3.2.3. Analysis over time

To characterize the time response of the adhesions, the measurements depicted in Fig. 4 were applied to beads subjected to 3 trapping conditions: a strong trapping k = 0.26 pN/nm (15 beads), a medium trapping k = 0.13 pN/nm (9 beads) and no trapping (19 beads). Fig. 6 depicts the change in AA intensity normalized to the value at t = 1min (Fig. 6a) and the difference in FRET efficiency compared to t = 1min (Fig. 6b) as a function of time. Mean values and standard deviations for all the data shown in Fig. 6a and 6b can be found in supplementary Fig. S6 and individual bead traces vs. time may be found in Figs. S7 and S8.

**Figure 5:**
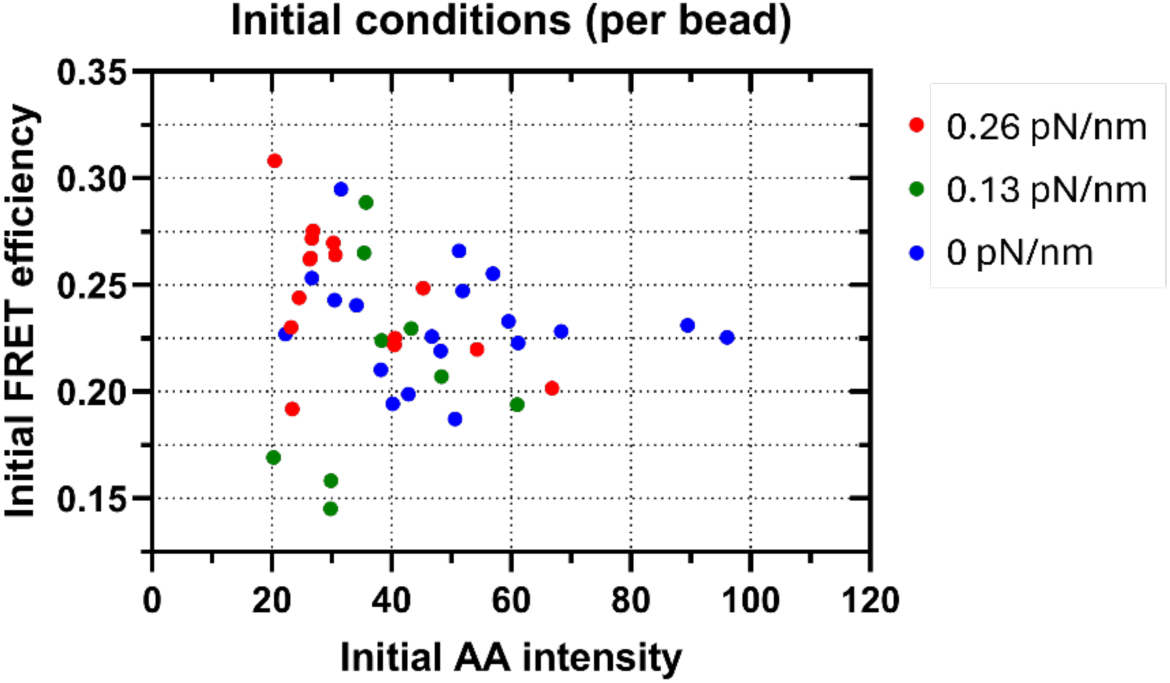
Initial FRET efficiencies vs AA intensities before turning on the laser trap. Each dot represents the mean value of the focal adhesions around a given bead. Colors indicate the laser trap stiffness that will later on be applied on each bead, Blue: no laser, Green: 0.13 pN/nm, Red: 0.26 pN/nm. For 0.26 pN/nm condition, data were obtained over 4 experimental days from 7 independent biological samples including measurements from 15 beads on 15 cells. For the 0.13 pN/nm condition, data were obtained from 2 independent biological samples prepared over 2 experimental days, comprising measurements from 9 beads on 9 cells. For the 0 pN/nm condition, data were obtained from 5 independent biological samples prepared over 3 experimental days, comprising measurements from 19 beads on 12 cells.

**Figure 6:**
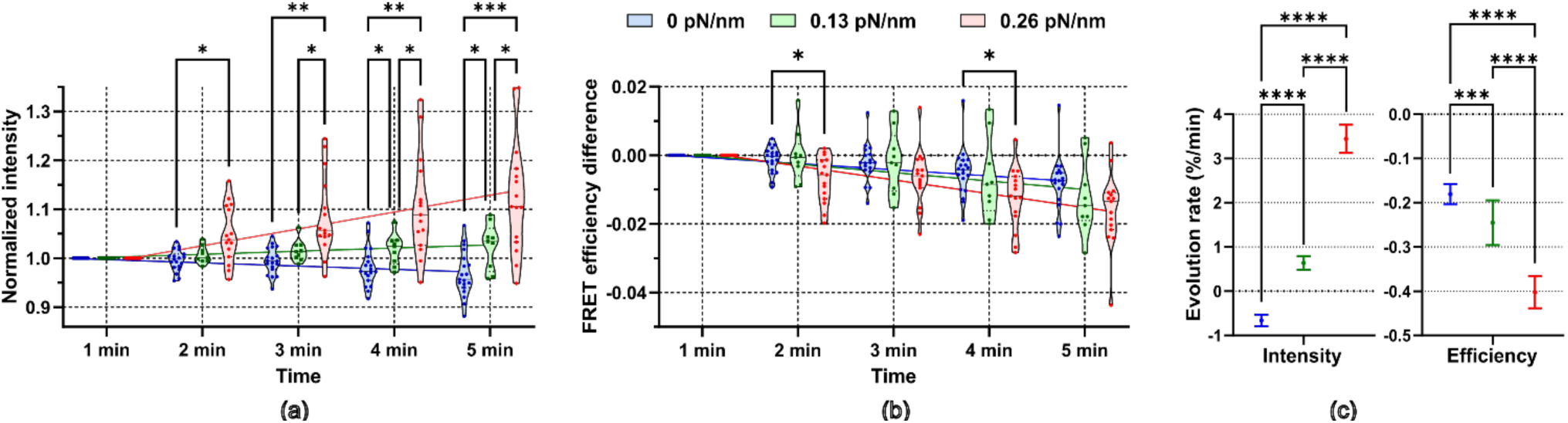
Violin plots of AA intensities normalized to t= 1min (a) and of FRET efficiency differences compared to t= 1min (b) as a function of time. Each point represents the mean value of the focal adhesions around a given bead. The data are reported for beads with no trapping (blue points), medium trapping (green points, k=0.13 pN/nm) or strong trapping (red points, k=0.26 pN/nm). t= 1min represents 1 min after the trapping laser is turned on. The lines are linear regressions of intensities and efficiencies versus time with a constraint to pass through the point (1,1) for intensity or (1,0) for FRET efficiency. Significant differences between the different groups are denoted by * (p<0.05), ** (p< 0.01), and *** (p<0.001) (Two-way ANOVA with repeated measures followed by Tukey’s multiple comparisons test) (c) Intensity slopes obtained from the linear regressions are respectively (% change/min) −0.66 ± 0.13, 0.64 ± 0.16 and 3.4 ± 0.32 for the different trap stiffnesses; the slopes of the FRET efficiency differences are respectively (difference in % absolute FRET efficiency/min) −0.18 ± 0.02, −0.25 ± 0.05 and −0.40 ± 0.04. All 6 slopes were significantly different from zero (p<0.001). Significant differences between the slopes are denoted by *** (p<0.001) and **** (p<0.0001) (One-way ANOVA followed by Tukey’s multiple comparisons). Number of repeats is the same as in Fig.5. The results of the statistical tests in Panels (a)-(c) are tabulated in supplementary Tables S1 – S7.

Comparisons between the different trapping conditions show statistically significant differences at each time point for relative intensities but not for FRET efficiency differences. To take into account the evolution over the 4 time points, a simple linear regression was applied to the data for each of the trapping conditions yielding the fitted slopes shown in Fig. 6c. Statistical analysis of the slopes, which takes into account the evolution as a function of time, reveals a significant difference between the different trap stiffnesses.

VinTS intensity increases with time and with trap stiffness while the intensity decreases by a few percent in the absence of trapping (the slope is slightly negative). The rate of AA intensity increase is also higher at 0.26 pN/nm compared with 0.13 pN/nm (Fig. 6c).

On the other hand, the FRET efficiency decreases in all 3 trapping conditions with higher rates of decrease as a function of trap stiffness (Fig. 6c). However, despite this observed increase in the magnitude of the slopes as a function of trap stiffness, the rates of FRET efficiency decrease were low compared to the rates of intensity increase. Also, the final FRET efficiency difference obtained at t= 5 min for both trap stiffnesses (decrease by 1-2% in absolute efficiency or ∼10% change relative to initial conditions) was not significantly different from that in the absence of the laser trap (Fig. 6b t= 5min). Thus, the adhesion’s response to the increased trap stiffness is dominated by an increase in vinculin recruitment rather than an increase in vinculin tension. The FRET efficiency difference measured at t = 5 min did not depend on the initial FRET efficiency (Supplementary Fig. S9).

#### 3.2.4. Intensity increase and FRET efficiency difference correlate at higher stiffness

To further elucidate the effect of trap stiffness and investigate the relationship between vinculin recruitment and tension, we plotted the AA normalized intensity as a function of FRET efficiency difference (Fig. 7) at all times between 1 and 5 min, with color indicating the trap stiffness. This plot is separated in four quadrants corresponding to different trends in the evolution of focal adhesions. In the upper right quadrant, intensity and FRET efficiency increase, which correspond to more vinculin recruited with less tension on each one, a trend that has been associated in the literature with assembling adhesions (44). In the lower right quadrant, vinculin intensity decreases while tension increases, a behavior associated with disassembling adhesions. The lower left quadrant where both intensity and FRET efficiency decrease and where most data points with no laser are located, could be interpreted as an effect of photobleaching by the LED. However, the decrease in AA intensity observed for VinTS (Fig. 7 lower-left quadrant) is lower than the decrease in AA intensity expected in a 12s exposure based on the photobleaching rate of diffuse mVenus in the cytoplasm (t1/2= 40s Fig. S1). Finally, in the upper left quadrant, where most of our data points at higher stiffness are located, vinculin intensity increases while FRET efficiency decreases, corresponding to more vinculin accompanied by higher vinculin tension. In addition to this trend, these data reveal a significant correlation between the increase in vinculin intensity and the decrease in FRET efficiency at 0.26 pN/nm (linear regression R2=0.38, p<0.0001) compared with no correlation at 0.13 pN/nm (R2=0.07, p=0.08) or with no laser trap (R2=0.002, p=0.65).

**Figure 7:**
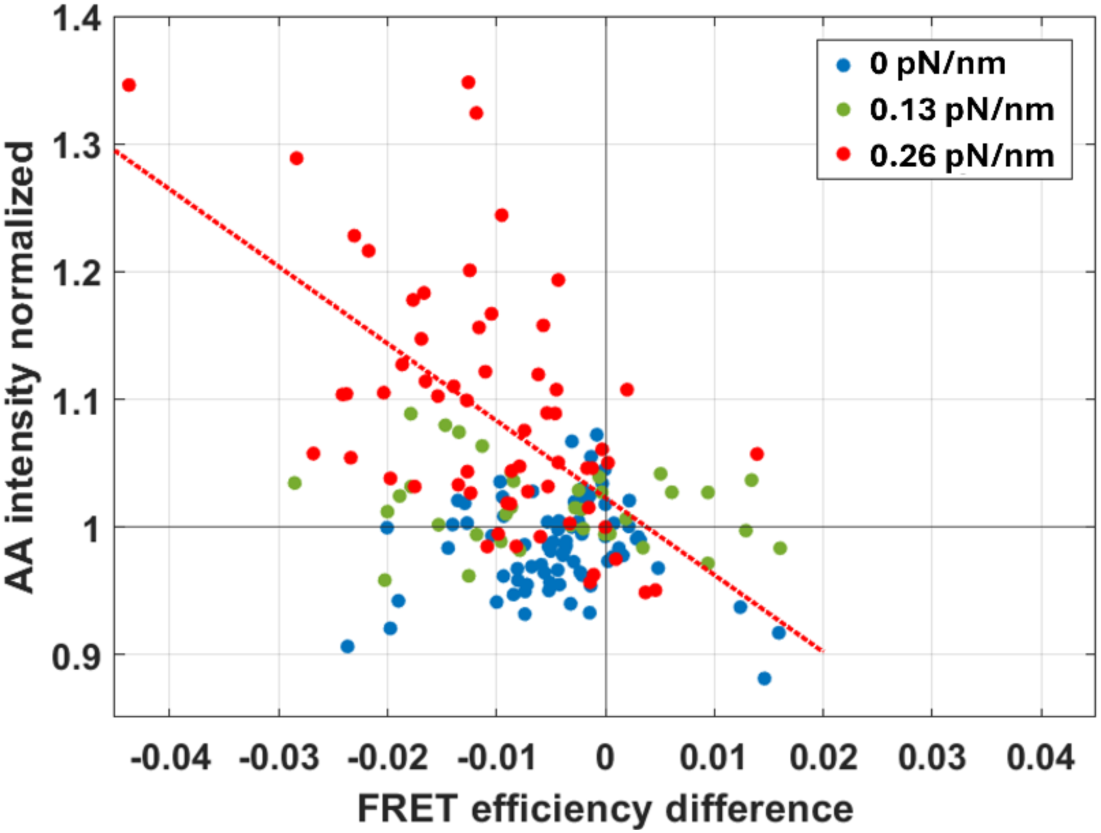
Normalized AA intensities of the focal adhesions around each bead as function of FRET efficiency difference plotted for all time points. Blue points are the beads without laser trapping, green are medium trap stiffness and red high trap stiffness. Black lines indicate the areas where normalized intensity increases (above 1) and FRET decreases (left of 0). The red dotted line is the linear regression among the red points subjected to higher trap stiffness (R^2^ = 0.38, slope significantly non-zero p-value<0.0001). Number of repeats is the same as in Fig.5. A measurement is taken each minute for 5 min, each bead gives 5 measurements.

The possible effect of photobleaching attributed to the lower left quadrant of Fig. 7, is difficult to gauge since vinculin turnover can compensate for the effect of photobleaching on vinculin intensity. While the effect of photobleaching can be mitigated by judiciously setting the LED excitation power levels based on the half-life of the fluorophores, photobleaching cannot be completely eliminated. Still, photobleaching alone cannot explain the differences in the results observed at the different trap stiffnesses, as the same LED power and exposure time were maintained in all these experiments.

### 3.3. Intensity and FRET efficiency as a function of laser force

The bead displacement at 5min (Fig. 8a) was used to calculate the force experienced by the bead at 5min using the calibrated trap stiffness. This analysis was used to investigate the relationship between vinculin intensity and vinculin tension as a function of force at the 5 min timepoint. The median displacement remained around 200nm (corresponding to a speed of ∼40 nm/min) whether there was no laser trap (207 ± 161 nm), medium trap stiffness (178 ± 338 nm) or high trap stiffness (199 ± 173 nm). The measured forces experienced by individual beads at t= 5 min vary between 8 pN and 90 pN when the optical trap is on, with the larger forces (>50pN) reached at the higher trap stiffness. However, the largest forces experienced at t= 5 min are not necessarily associated with the largest increases in vinculin recruitment, and relative increases in vinculin intensity above 1.1 could be observed at high laser trap stiffness but at moderate forces between 20 pN and 30 pN (Fig. 8b). By varying the stiffness of the optical trap, the results show that the increased vinculin recruitment is governed by the stiffness of the trap (red points vs green points in Fig. 8b) rather than the magnitude of the traction force. By comparison, the decrease in FRET efficiency observed at t= 5 min, and thus the vinculin tension, does not appear to be correlated with the force reached at 5 min nor with the trap stiffness (Fig. 8c). Here we plotted the FRET efficiency normalized to t=1min rather than the FRET efficiency difference as in Fig. 6b to show that the relative change in intensity can be up to 35% while the relative change in FRET efficiency is at most 15%. Individual bead traces of normalized AA intensity and FRET efficiency difference vs. force may be found in supplementary Fig. S10.

**Figure 8:**
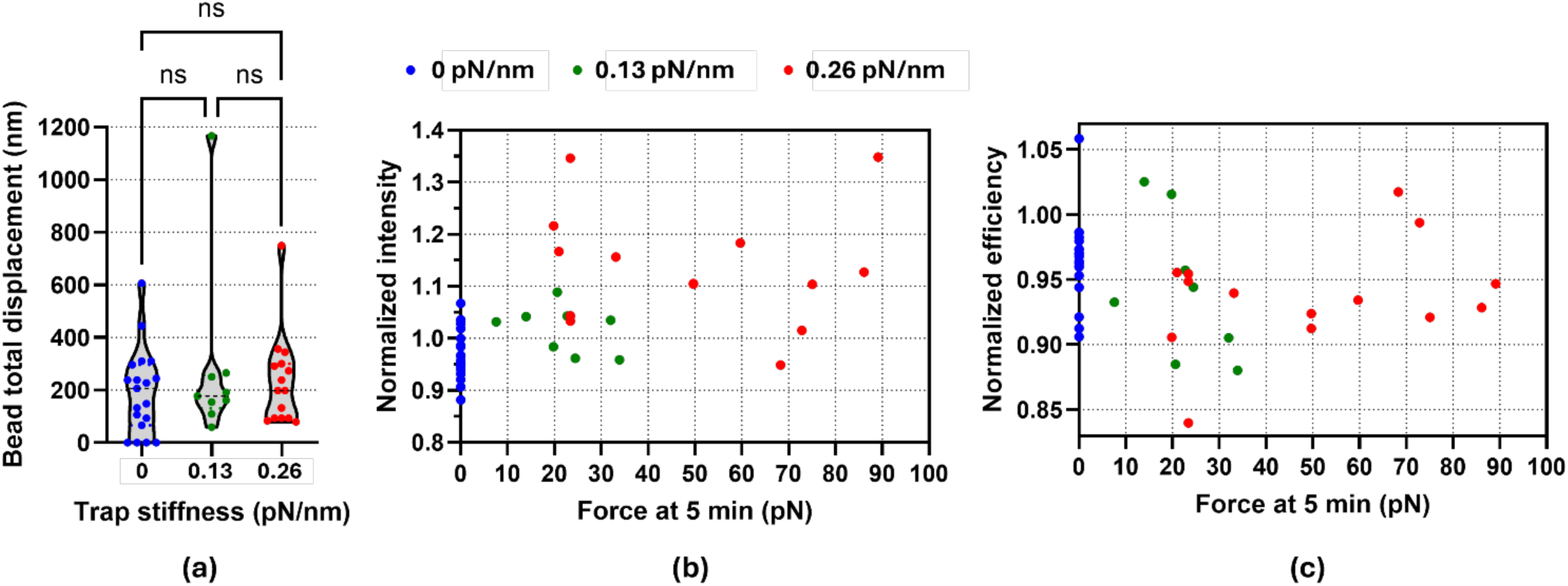
(a) Displacement of the beads from their initial positions to their positions at time t=5 min for different trap stiffnesses. No significant difference was found between the different groups (Ordinary one-way ANOVA followed by Tukey’s multiple comparisons test). The results of the statistical tests in (a) are tabulated in supplementary Table S8. (b) and (c) Normalized AA intensities and FRET efficiencies of the focal adhesions around each bead as function of the force applied by the laser on the bead after 5 min. Blue points are the beads without laser trapping, green are medium trap stiffness and red high trap stiffness. Number of repeats is the same as in Fig.5. However, the beads displaced by more than 500 nm are not shown on force plots on (b) and (c) since they are not in the linear domain of the trap, (b) and (c) shows 14 beads for 0.26 pN/nm, 8 beads for 0.13 pN/nm and 19 beads for 0 pN/nm.

### 3.4. Focal adhesions moving away from beads

The vinculin puncta which formed around the beads typically remained on the surface of the bead and moved together with the beads at a speed of ∼40 nm/min (Fig. 8a). In addition, the focal adhesions on a given bead experienced similar changes in vinculin intensity or FRET efficiency, justifying their averaging for a given bead (data in Figs. 6-8). However, a few focal adhesions behaved very differently. An example is shown in Fig. 9 (and supplementary movie), where we recorded the FRET images over 11min. In this case, the bead moves to the left, in the negative x-direction at 46nm/min and the two vinculin puncta on the left of the bead (numbered 1 and 4, Fig. 9c) move to the left with the bead and remain on the bead’s surface. However, the vinculin puncta on the right (numbered 2 and 3, Fig. 9c) move to the right and away from the bead’s surface at higher speeds of 177 nm/min (adhesion 2) and 185 nm/min (adhesion 3). Vinculin intensity for adhesions 1 and 4 remains constant for 7min but slightly decreases over the 11min. On the contrary, intensity of adhesions 2 and 3 increases up to 7min and then remains constant. The vinculin FRET efficiency of adhesions 1 and 4 starts at about 0.28 and decreases by 0.02 over the first 5 minutes then remains constant except for a brief jump at t=10min, while the FRET efficiency of the faster moving adhesions 2 and 3 continues to decrease as they move away from the bead, suggesting that vinculin is experiencing higher tension as it is getting pulled to the right. We observed such vinculin puncta moving away from the bead’s surface in 3 other beads tested at 0.26 pN/nm over 5 min.

**Figure 9:**
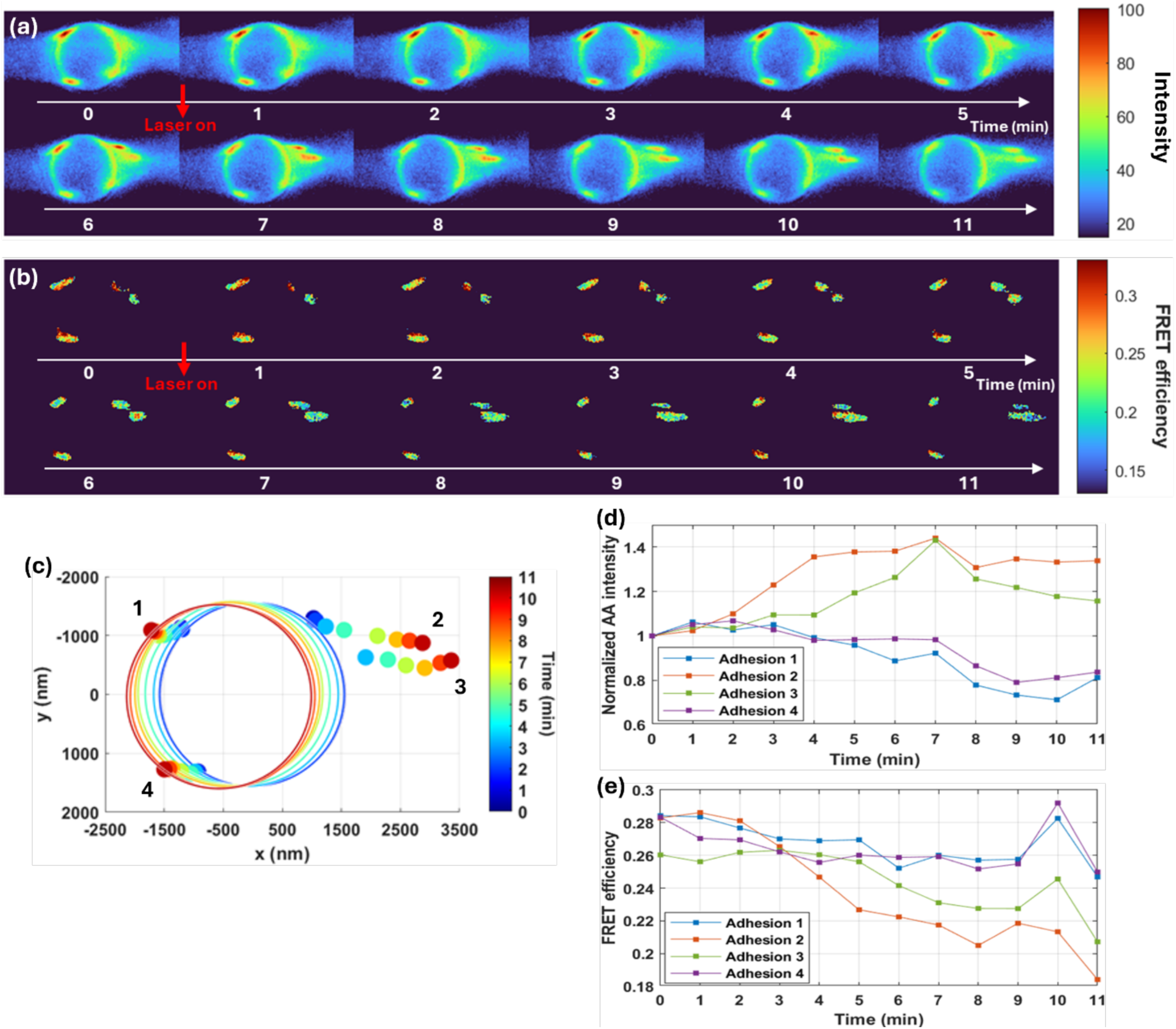
Example of focal adhesions moving away from a bead trapped with high stiffness (0.26 pN/nm). (a) and (b) are respectively total intensity (sum AA+DD+DA) and FRET efficiency images of the focal adhesions around the bead. The procedure described in Fig. 4 was used but images were recorded over 11 min. (c) Successive positions of the bead (large circles) and of the four focal adhesions (dots 1, 2, 3, 4). (d) and (e) are the normalized AA intensity and absolute FRET efficiency of each focal adhesion vs time. The bead and two adhesions (1 and 4) move to the left at a speed of 46 nm/min while adhesions 2 and 3 are moving to the right at 177 nm/min (adhesion 2) and 185 nm/min (adhesion 3).

## 4. Discussion

To quantify the response of vinculin to a local mechanical stimulus, we combined optical tweezers and FRET efficiency measurements of the vinculin tension sensor, VinTS. By functionalizing the surface of polystyrene beads with fibronectin, which can attach to integrins at the cell surface, we measured the traction force imparted to the beads by the cell and the beads displacement while quantifying vinculin recruitment and tension at the focal adhesions formed on the beads’ surface. The direction of the bead’s displacement and of its associated force is controlled by the cell. The presence of the optical trap on a bead counteracts the bead’s movement and allows us to measure the cell force as a function of increasing laser power (optical trap stiffness). Since the cell is pulling the bead rather than migrating on a substrate, our experiments may be more akin to matrix remodeling (57) rather than to cell migration, as the adhesions we are considering are formed with the bead on the dorsal surface of the cell.

Beads, 3µm in diameter, were tested with trap stiffnesses of 0.26 pN/nm or 0.13 pN/nm or with no trapping laser. Over the course of 5 min, the median bead displacement was on the order of 200nm and did not depend on trap stiffness while forces reached 8-35pN at 0.13 pN/nm and 20-90pN at 0.26 pN/nm (Fig. 7). At the lower 0.13 pN/nm trap stiffness, the adhesion response involved an increase in vinculin tension accompanied by a moderate increase in vinculin recruitment (<10% intensity increase) while at 0.26 pN/nm, a similar increase in vinculin tension was accompanied by a significantly larger increase in vinculin recruitment (up to 35% intensity increase). In the absence of the laser trap, the decrease in FRET efficiency was accompanied by an average slight decrease (<5%) in vinculin intensity. Thus, the response to increased trap stiffness was governed by increased recruitment of vinculin rather than further increase of vinculin tension. The external force applied on the trapped bead is distinct from the force across the vinculin molecules. The force on the bead reports on the cell traction force which is effected through multiple proteins acting at the adhesion site. One goal of this study was to investigate how this traction force translates into tension across vinculin. The FRET efficiency of the vinculin tension sensor reports on the mean tension across an individual vinculin protein within an imaged voxel. As such, our data show that as the cell traction force increases from a few up to 90pN on the bead, VinTS FRET efficiency decreases by 1-2% (in absolute FRET), which corresponds to ∼0.5-1pN increase in tension across vinculin based on the force-calibrated tension module TSMod inserted in the VinTS probe (44).

The increased vinculin recruitment observed at higher stiffness could not be fully accounted for by the larger magnitude of the cell forces reached at t= 5 min since both high (>50pN) and moderate (20-30 pN) forces could be associated with high vinculin recruitment (>1.1) (Fig. 7b). Instead, our data revealed a correlation between the increase in vinculin intensity and the decrease in FRET efficiency at 0.26 pN/nm that was not observed at 0.13 pN/nm or without laser (Fig. 8). This behavior is different from the relationship between vinculin intensity and tension observed as a function of time during focal adhesion assembly or disassembly (44). Indeed, during assembly, the total vinculin signal increases while vinculin tension starts high and subsequently decreases, and during disassembly, the total vinculin signal decreases while vinculin tension remains low or further decreases. Static measurements of vinculin intensity, such as the initial values reported in Fig. 5, were not correlated with FRET efficiency, corroborating published data showing that VinTS intensity is generally not correlated with FRET efficiency at focal adhesions in various states of assembly or disassembly (49). In this context, our experiments, which are conducted on adhesions in different initial states, demonstrate that a positive correlation between vinculin recruitment and vinculin tension can be induced by the applied external laser force over time. Further study of adhesion dynamics is required to elucidate the mechanisms underlying the interplay between vinculin tension and recruitment in response to sustained external forces.

In the case of 4 beads tested at 0.26 pN/nm, we observed vinculin puncta moving away from the trapped bead. An example in which the movement was in the direction opposite to the bead’s displacement was shown in Fig. 9. The moving puncta exhibited both an increase in vinculin intensity and an increase in vinculin tension (decrease in FRET efficiency) (Fig. 9). This suggests that we are not observing a focal adhesion disassembly with disengagement of vinculin. Moving focal adhesions have been previously observed during fibronectin assembly (58,59). The reported speed of those adhesions driven by actin stress fibers (58) was 6.4µm/h, which is approximately half the movement speed we observed in Fig. 9 (∼0.18µm/min, which gives ∼11µm/h). Future studies with additional labels for paxillin, actin, fibronectin and other components of the focal adhesion complex are required to fully interpret the mechanism leading to the flow of vinculin away from the bead in response to force.

Adhesion maturation and reinforcement was previously observed on fibronectin functionalized beads attached to cells and placed in an optical trap as evidenced by adhesion growth and vinculin recruitment (24–26). The forces and bead displacements measured in this study are comparable to reported values for fibronectin-coated 4.5 µm beads in an optical trap with 0.16 pN/nm stiffness (24,26). However, in other experiments on mouse embryonic fibroblasts involving smaller beads, displacements of ∼100nm/s were reported for fibronectin-coated 1µm beads in a 0.4pN/nm optical trap (25) and actin flow without the addition of functionalized beads is typically on the order of 5nm to several tens of nanometers per second at focal adhesions formed on a rigid substrate (13,42,43,60,61). Speeds of ∼3µm/min (62) or ∼0.5µm/min (63) were also reported in early studies on the retrograde movement of 0.2µm-1µm sized particles on the membrane of living cells. Assuming the bead movement is governed by actin flow, which is supported by the detection of actin fibers ending at the adhesion complexes forming around the beads (Fig. 3) and previous literature (5), the comparatively low speed of bead displacement measured for the 3µm beads in our study, or the reported 4.5 µm beads (24,26), suggests that actin flow speed may be slowed by the larger bead size. Furthermore, despite the increase in vinculin recruitment, the bead displacement in our data did not change as a function of laser trap stiffness as might be expected from the molecular clutch model (5,12,13,20,46). Notably, our data corroborate previous results showing no change in bead velocity as a function of trap stiffness in accordance with a feedback mechanism balancing the recruitment of acto-myosin motors with increasing traction force (29). This may be a consequence of the slow bead displacement and the loading rates involved in our experiments placing our conditions in the low-stiffness range of the clutch model where actin retrograde flow may remain constant until a higher stiffness is achieved (64).

Limitations of this study that can be addressed in future work include the use of VinTS mutants to improve biological interpretation of our data and identify the vinculin functions and molecular mechanisms at play. The tail-less vinculin tension sensor (VinTL) (44) or the mutant VinTS-I997A (50), which cannot bind actin and remain unloaded, can be used to establish the unloaded, FRET efficiency baseline under our experimental conditions and confirm that the 1-2% decrease in FRET (Fig. 6b) is lower than this baseline. Furthermore, the unloaded control can be used to investigate if vinculin recruitment requires vinculin tension and if the correlation between vinculin recruitment and vinculin tension (Fig. 7) is causal. Mutants of the vinculin tension probe specifically altering the directional catch bond between vinculin and actin (65) could provide additional insight as could mutations affecting the autoinhibitory interaction between vinculin’s head and tail domains (66–68) or dimerization of the vinculin tail which allows actin bundling (69). Labeling other proteins such as actin, talin or paxillin, as well as fibronectin on the bead’s surface may help explain the movement of adhesions away from the bead (Fig. 9). While the involvement of integrins was not directly demonstrated in this study, this could be achieved by testing cells that are treated with anti-Beta1 integrin antibody before adding the FN-coated beads or by comparing Beta-1 integrin antibody coated beads to FN-coated beads. Finally, modulating myosin II-dependent contractility in conjunction with our measurements would help put our work in the context of previous studies on the role of actin stress fibers (34,42) and acto-myosin contractility (14,23) in focal adhesion maturation.

In conclusion, by combining optical tweezers with the vinculin tension sensor we investigated the role of vinculin tension at cell adhesions in response to force and optical trap stiffness. Our results show that the adhesion’s response to optical trap stiffness is dominated by an increase in vinculin recruitment while bead displacement and vinculin tension are maintained and motivate additional inquiry into the mechanisms by which forces control the interplay between vinculin recruitment and tension.

## Data availability

The raw data supporting the conclusions of this article and the code used for data analysis in this publication will be shared upon reasonable request to the authors.

## Author Contributions

C.D. performed the experiments and analyzed the data both in Palaiseau and in Rutgers. R.I.C. prepared the lentivirus and the transduced fibroblasts expressing VinTS. N.N.B., and N.W. assisted with data analysis. C.D., R.I.C., N.N.B. and N.W. wrote the paper.

## Declaration of interests

C.D., NW. and N.N.B. declare no competing interests. R.I.C. is currently employed at The Well Bioscience Inc.

## Acknowledgements

We thank Prof. Marie Erard (Univ Paris-Saclay) for help in transfection of the fibroblast cells with VinTS before we could create the lentivirus transduced cells. We thank Prof. Brenton Hoffman (Duke University) for the pRRL-VinTS plasmid and for pertinent discussions. CD was supported by a PhD grant from the French Ministry of Higher Education through the Ecole Doctorale EDOM, plus financial support from Laboratoire Charles Fabry for her missions at Rutgers. This work was supported by Centre National de la Recherche Scientifique (CNRS) through a Tremplin@INP2020 grant, by a grant “Investissements d’Avenir” of the LabEx PALM (N° ANR-10-LABX-0039-PALM) and by a grant part of the France 2030 program (N° ANR-11-IDEX-0003) to N.W. and by NSF award CMMI-1825433 and a Rutgers Global International Collaborative Research Grant to N.N.B.

## Supplementary Material

- One document including Supplemental Methods, Fig. S1-S10 and Tables S1-S8.
- One supplemental movie (animated version of Fig.9)

## Supporting Citations

References (70–75) appear in the supplemental material and methods.

## Notes

### Summary of Updates

Abstract shorted; Novelty more clearly stated in Introduction and Discussion; Data on individual beads and statistical tests added in supplemental files

