## Supplemental material for "Optical tweezers combined with FRET tension sensor reveal force-dependent vinculin dynamics"

### Supplemental material and methods
### Optical tweezers combined with FRET tension sensor reveal force-dependent vinculin dynamics

**Camille Dubois,[1] Rick I. Cohen,[2,3] Nada N. Boustany,[2,**] Nathalie Westbrook [1,*]**
*1 Université Paris-Saclay, Institut d'Optique Graduate School, CNRS, Laboratoire Charles Fabry, Palaiseau, France*
*2 Rutgers University, Department of Biomedical Engineering, Piscataway, New Jersey, United States*
*3 Present address: TheWell Bioscience Inc., Monmouth Junction, New Jersey, United States*

#### Supplemental methods: lentivirus packaging

On Day 0 HEK293FT cells were passaged from T75 flasks using TrypLE, counted using Trypan Blue exclusion dye, and plated at a density of 72,000 cells/cm$^2$ onto tissue culture treated 10 cm dishes in DMEM/F12 containing 10% FBS and 10 µg/mL gentamicin medium. The following day (Day 1), the medium was removed, the plate gently rinsed once with warm DMEM/F12 base media (contains 2 mM glutamax, 15 mM HEPES, and 1x non-essential amino acids) with no serum or antibiotics and replaced with 7 mL of warmed Optimem containing 2% of heat-inactivated FBS. The plates were placed back into the incubator while the transfection mixtures were prepared. For each 10 cm dish transfected, plasmids (a total of 20 µg of DNA) were diluted into a final volume of 930 µL of Optimem in a sterile 2 mL low DNA binding centrifuge tube in the following amounts; 10 µg of pRRL-VinTS (gift from Brenton Hoffman); 6.67 µg of psPAX2 (gift from Didier Trono (Addgene plasmid # 12260 ; http://n2t.net/addgene:12260 ; RRID:Addgene_12260); and 3.3 µg of pMD2.G (gift from Didier Trono (Addgene plasmid # 12259 ; http://n2t.net/addgene:12259 ; RRID:Addgene_12259)). The DNA was mixed well by inverting the tube several times and then centrifuged. A 50 kDa PEI Solution (Sigma) was prepared according to Boussif and coworkers (1) and used at a ratio of 3.5 µL of PEI per µg DNA; 70 µL was added to each tube with 930 µL DNA/Optimem mixture, capped, and mixed by rapid shaking by hand for 10 seconds. The tubes were centrifuged and set aside for 15 min incubation for PEI/DNA complexing. To ensure even transfection, the mixture was added dropwise to each 10 cm dish of HEK293FT cells while gently swirling the media and moving pipette tip. The dishes were replaced into the incubator until the following day. On Day 2, the dishes were inspected for viability and fluorescence caused by the expression of plasmid. The medium was changed to 10 mL of warmed DMEM/F12 (as above) with 10% of heat-inactivated FBS, 10 µg/mL gentamicin and 4 mM caffeine (2). On Day 3, the medium was collected and replaced with the same without caffeine. On day 4 the medium was collected, and the dishes discarded. The collected supernatant on each day was centrifuged at 4$^o$C at 500 x g in 50 mL tubes in vapor-proof centrifuge carriers. The clarified supernatant (up to 35 mL) was filtered using sterile 0.45 µm PES tube top filters (Corning), and transferred into 38.5 mL tubes (Beckman Coulter, 326823) held in 4$^o$C cooled ethanol and UV light sterilized SW-28 ultra-centrifuge holders (Beckman Coulter). Each holder was balanced using a scale and sterile DMEM/F12 base medium. The supernatant was centrifuged for 2 hours at 25,000 RPM (average calculated g force of 82,700 x g), removed and pellet resuspended in 100x less volume (i.e. 350 µL) of PBS-EDTA with 5% Trehalose.

#### Photobleaching measurements of mTFP1 and mVenus

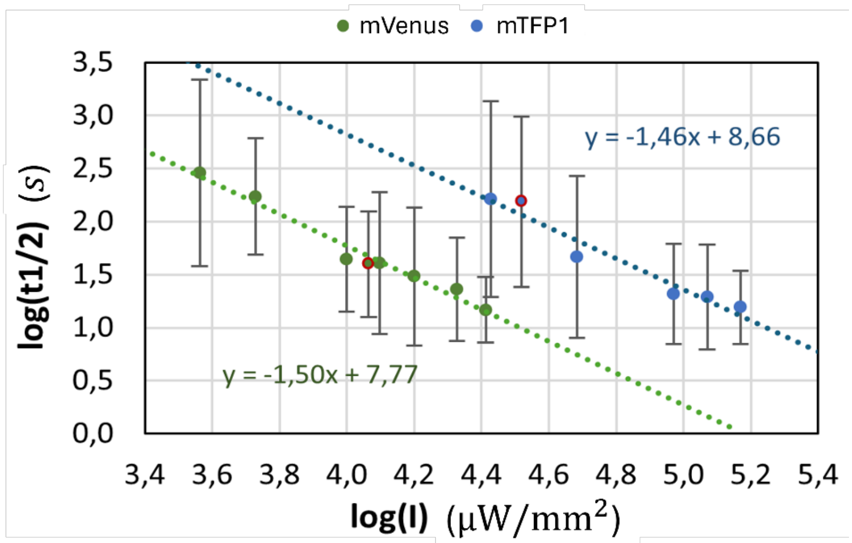

***Figure S1 :*** *Photobleaching measurements of mTFP1 and mVenus. Log-log plot showing the effect of excitation intensity on the half life of each fluorophore measured on CHO-K1 cells expressing only mTFP1 or only mVenus. Points are measurements ± standard deviation and the dotted lines are linear regressions showing the supralinear dependency between the half life of the fluorophores and the excitation intensity* (3). *The red circled dots are the settings used for the measurements with* $11.6 \cdot 10^{3} \mu W/mm^{2}$ *for mVenus excitation (LED A) and* $33.0 \cdot 10^{3} \mu W/mm^{2}$ *for mTFP1 excitation (LED D).*

#### Optical tweezer calibration

##### 1) Lateral position of the bead

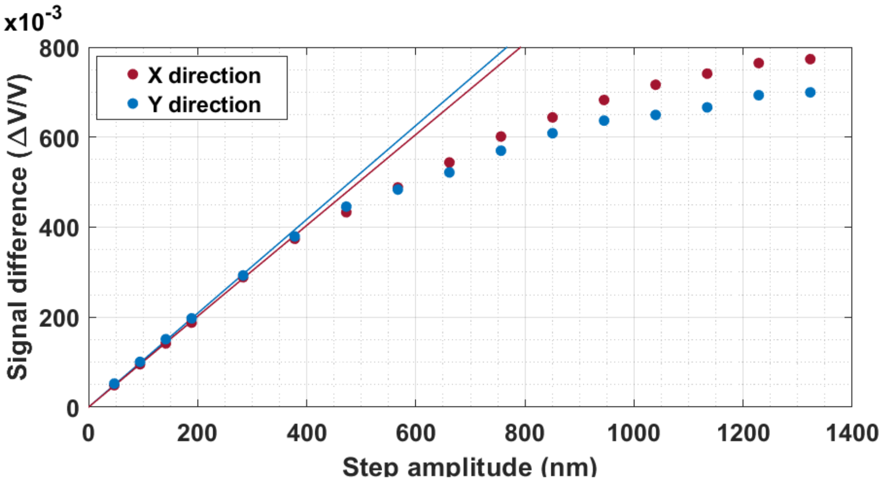

***Figure S2 :*** *Calibration of the displacement of the bead from the QPD relative voltage. Relative signal difference of voltage in X and Y directions in function of the distance of the bead from the trap center. The trap is quickly moved using the AOMs. The dotted lines are linear regressions on the experimental measurements until 400nm and give the conversions factors* $C_X = 1.00 \pm 0{,}02 \frac{\Delta V}{V} / \mu m$ *and* $C_Y = 1.02 \pm 0{,}03 \frac{\Delta V}{V} / \mu m$.

#### 2) Trap stiffness : details of the two calibration methods

- For the power spectrum density (PSD) analysis, a Brownian motion trajectory is acquired for 2 seconds at 62.5 kHz. The Fourier transform of the trajectory is fitted by a Lorentzian using the program published in (4,5). Fits are performed on frequencies from 50 and 5000 Hz because of the presence of low frequency noise (Fig. S3a). The trap stiffness is then given by : $k = f_c \times 2\pi\gamma$ with $f_c$ the trap cutoff frequency and $\gamma$ the friction coefficient.
- For the step-response method, an acquisition consists of at least 50 bead relaxations towards the center of the trap for a given trap displacement of 94.5 nm. Signals are averaged and fitted to obtain the relaxation times presented on Fig. S3b. The trap stiffness is deduced as $k = \frac{\gamma}{t_b}$, with $\gamma$ the friction coefficient and $t_b$the relaxation time.

The friction coefficient is given by the Stokes's law $\gamma = 6\pi\eta R$ with R the radius of the bead and $\eta$ the local viscosity. The local viscosity around the bead is corrected by Faxen's law to take into account the distance of the bead from the coverslip surface (4).

These two methods are compared using acquisitions on the same bead. The examples shown in figures S3a and b are acquired for a 3 µm bead at 370 mW at a distance $h = 5\ \mu m$ from the surface. The friction coefficient is then γ$(h = 5\ \mu m)$= 3.39.$10^{-8}$ kg.$s^{-1}$. For the example in Fig. S3a, the cutoff frequency in the x direction is 1208.4 Hz which gives a trap stiffness of $k = 0.258\ pN/nm$ (the same analysis in y direction gives $k = 0.241\ pN/nm$). One example of a step-response result is shown in Fig. S2b with exponential fits of the round trip of the bead for a step of 94.5 nm in the x direction. The average relation time is $t_b = 129.2\ \mu s$ which gives a trap stiffness of $k = 0.263\ pN/nm$ (the same measurement with a step in y direction gives $k = 0.261\ pN/nm$).

Fig. S3c show the stiffness of the trap as a function of laser power, obtained from the step-response method and the power spectrum analysis for 3 µm diameter beads. Stiffnesses obtained with the two methods are in close agreement and increase linearly with laser power. Fig S3d shows the trap stiffness for different bead diameters of 2, 3 and 6 µm. For bead diameters greater than the wavelength such as the ones we used, trap stiffness can be modeled by ray optics leading to a stiffer trap for smaller beads (6). For a laser power of 370 mW at the entrance of the objective (high stiffness), the trap stiffnesses measured using the step method and averaged in x and y are 0.68 pN/nm for 2 µm beads, 0.26 pN/nm for 3 µm beads and 0.22 pN/nm for 6 µm beads.

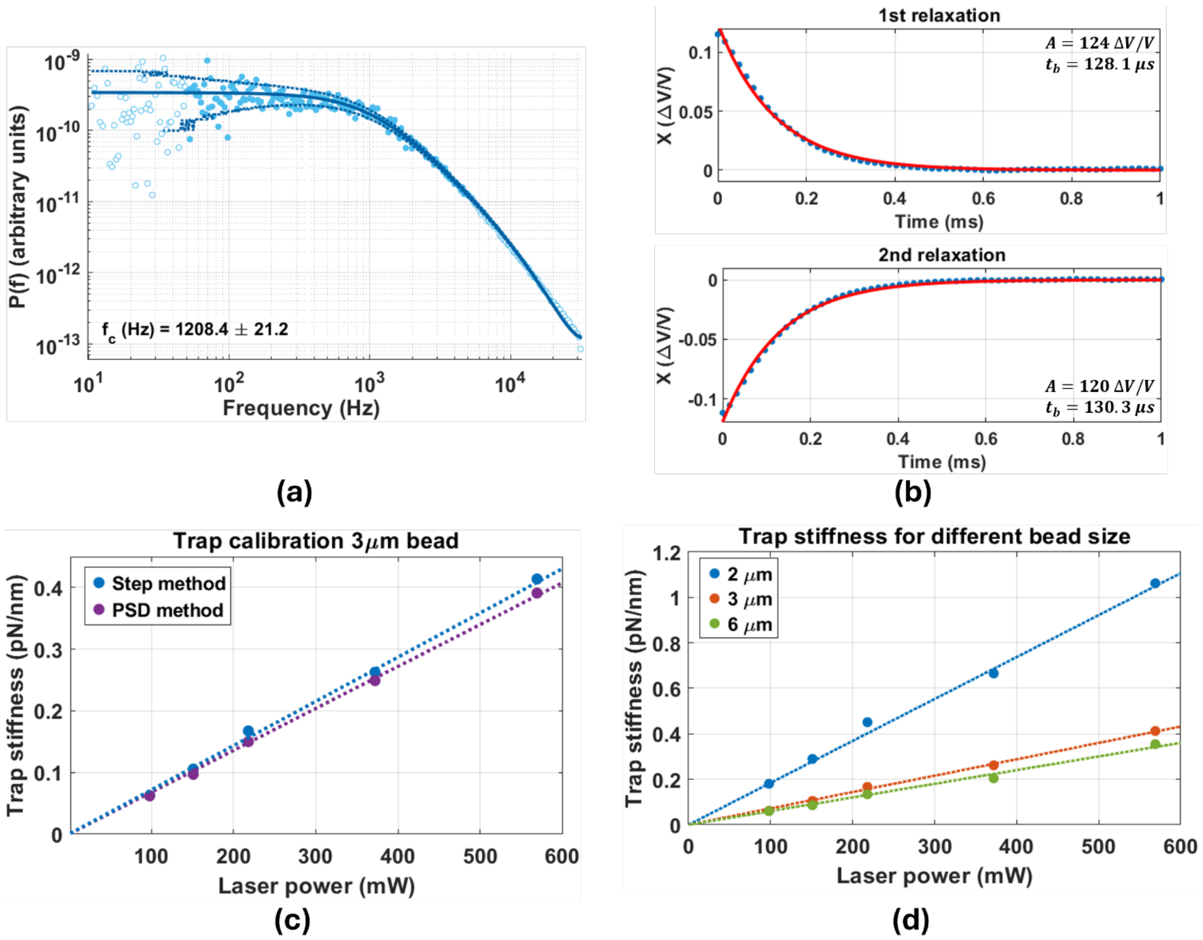

***Figure S3 :*** *Optical tweezer stiffness calibration. The same 3µm bead trapped in water at 5µm from the coverslip surface is used for the two methods of calibration. (a) Power spectral density of the Brownian motion acquired during 2s at 370 mW in the x direction. Each dot represents an average over 200 experimental data points. A Lorentzian fit is performed on frequencies between 50 and 5,000 Hz (solid dots). The solid line is the fit and the dotted lines indicate the uncertainty at ±the standard deviation. (b) Step-responses of a 3 µm bead at 370 mW for a step of 94.5 nm in the x direction. The two relaxations are the round trip of the bead averaged over 50 repetitions. The exponential fit* $A\,exp(-\frac{t}{t_b})$ *is given in red. (c) Trap stiffness for a 3µm bead obtained with the power spectrum analysis (in violet) and the step-response (in blue) methods as a function of the incident laser power measured at the entrance of the objective. Each point is an average of x and y directions. (d) Trap stiffness normalized by the laser power for bead diameters of 2, 3 and 6 µm using the step method. Linear regressions give the relation between stiffness and laser power: 1.87, 0.71 and 0.60 pN/nm/W.*

#### Confocal images of focal adhesions on several cells with different bead coatings

Fig. S4a shows more examples of fibroblast cells with FN coated beads in the same conditions as in Fig.3 of the main text, showing vinculin puncta (white arrows) connected to actin filament (blue arrows). We also tested other types of coatings for the beads. In the case of uncoated beads, the beads do not attach to the cells after 45 min of incubation and can be readily pulled away from the cell using the optical trap.  In the case of RGD coating, and after the 45 min incubation, the beads attach to the cell to the extent that they cannot be readily pulled away from the cell by the optical trap. In the confocal images (Fig. S4b), we observe a diffuse VinTS signal around the RGD coated beads similar to the diffuse signal within the cytoplasm and we do not observe actin organization around the bead.

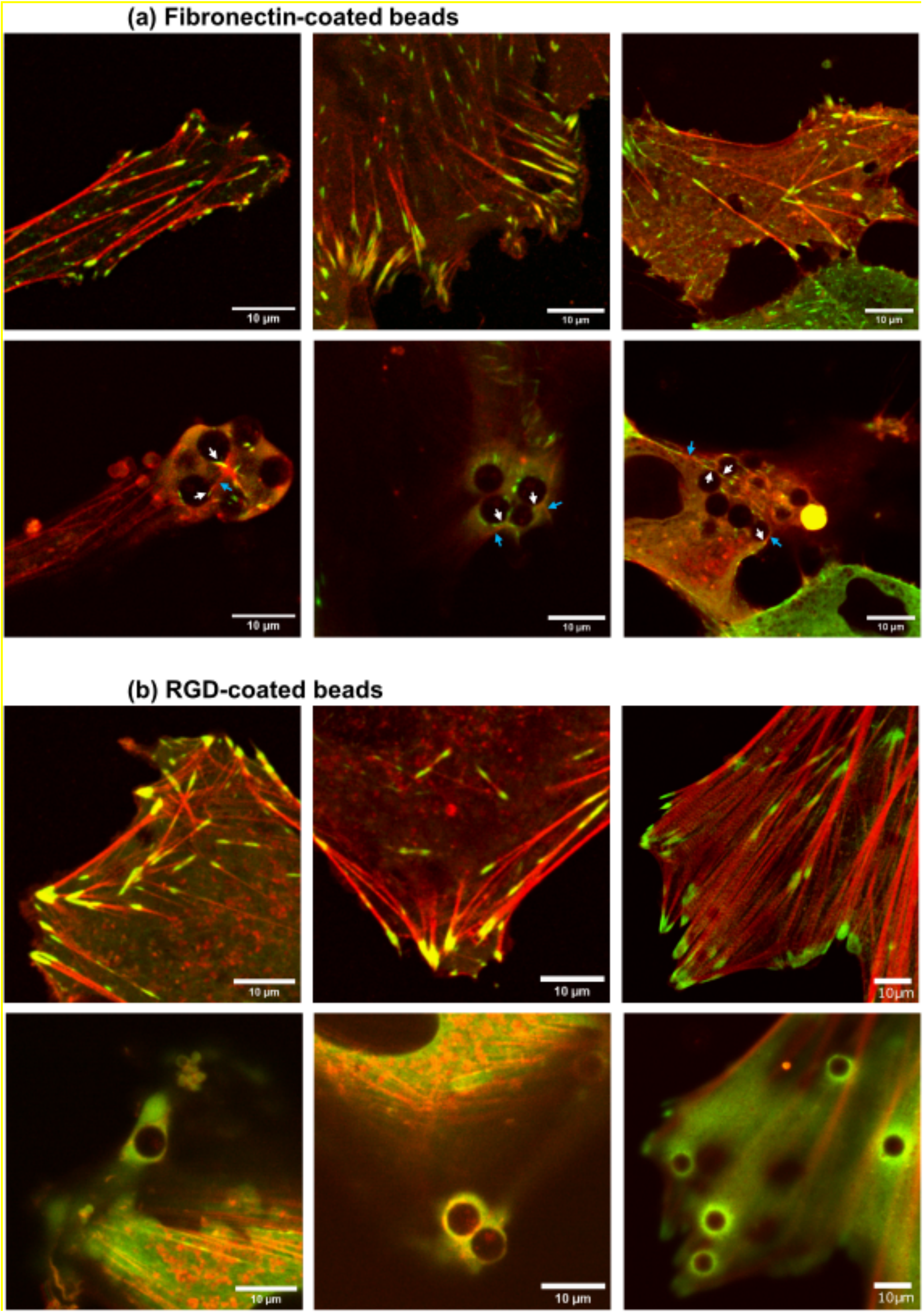

***Figure S4 :*** *Confocal images of fibroblast cells after 45 min of incubation with 6µm beads coated with (a) fibronectin or (b) RGD. Vinculin is marked using the tension sensor VinTS in green and actin using acti-RFP in red. Upper row shows the focal adhesions (FAs) on the coverslip and lower row is at the height of the beads. (a) Vinculin puncta are present on the surface of FN beads (white arrows on the lower row of images) and are linked to actin filaments (blue arrows). (b) RGD beads are also slightly embedded into the cells but the attachment is not specific since no vinculin puncta nor actin reorganization around the bead is observed.*

#### Intensity images under force as a function of time on several beads

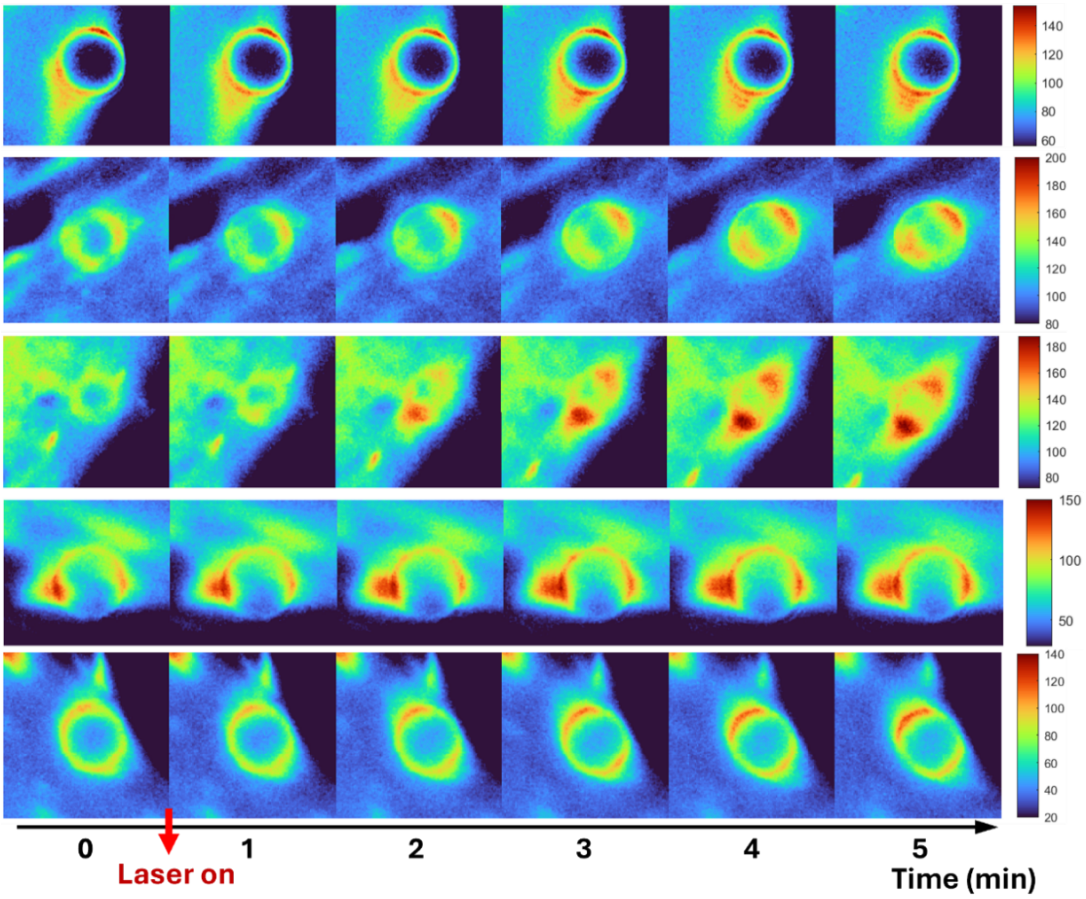

***Figure S5 :*** *Examples of intensity images of cells expressing VinTS when fibronectin coated 3µm-diameter beads are trapped with high stiffness (0.26 pN/nm). Fluorescence images are the sum of the three registered images DD, DA and AA recorded every minute. The laser is turned on immediately after the first image acquisition. The color scale is adjusted separately for each time series.*

### Vinculin intensity and FRET efficiency difference as a function of time: mean values and standard deviations

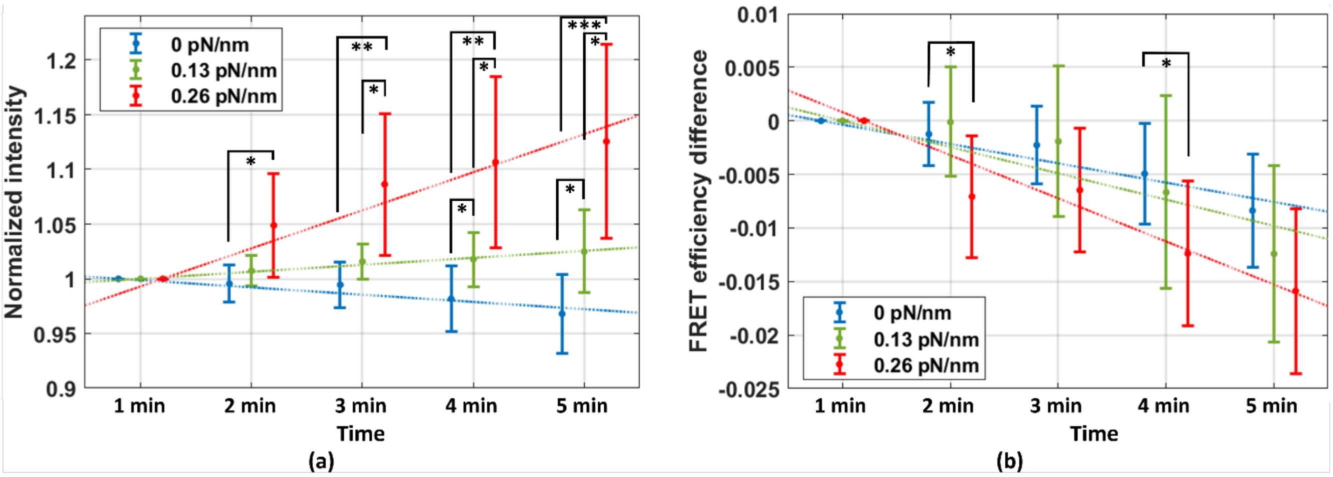

***Figure S6***: *Plots of AA intensities normalized to t= 1min (a) and of FRET efficiency differences compared to t= 1min (b) as a function of time. The data and conditions are the same as the violin plots in Fig.6 of the main text, but here we show the mean values ± standard deviation. The dotted lines are the same linear regressions as in Fig.6.*

### Vinculin intensity and FRET efficiency difference as a function of time for each bead:

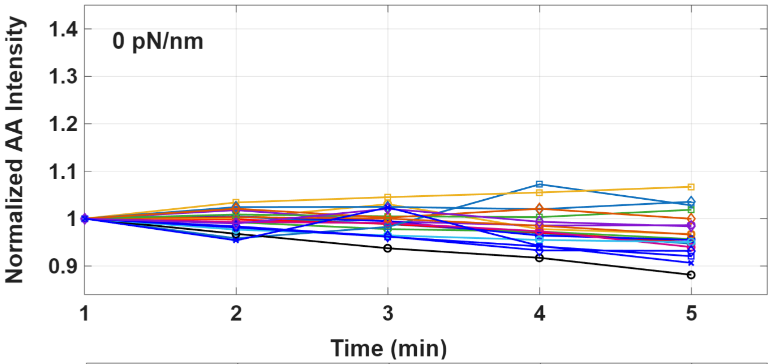

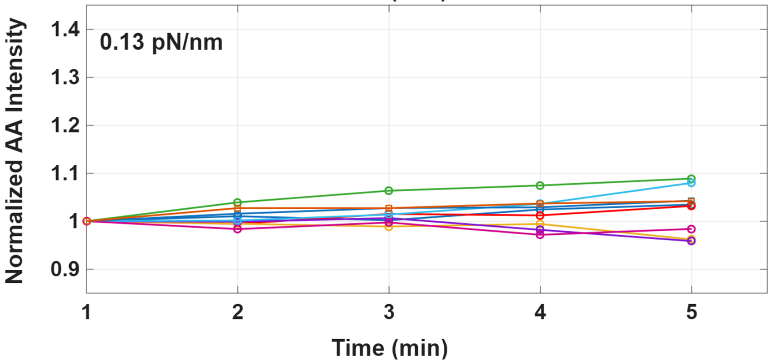

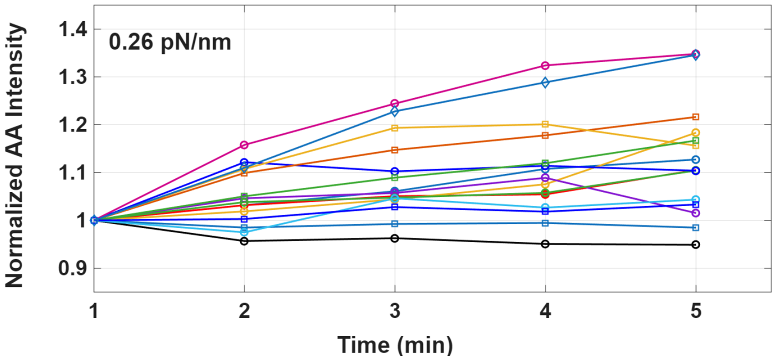

***Figure S7:*** *Plots of normalized AA intensity as a function of time for each bead at the 3 trap conditions (No trap (0pN/nm), 0.13pN/nm and 0.26 pN/nm). Each trace represents a different bead.*

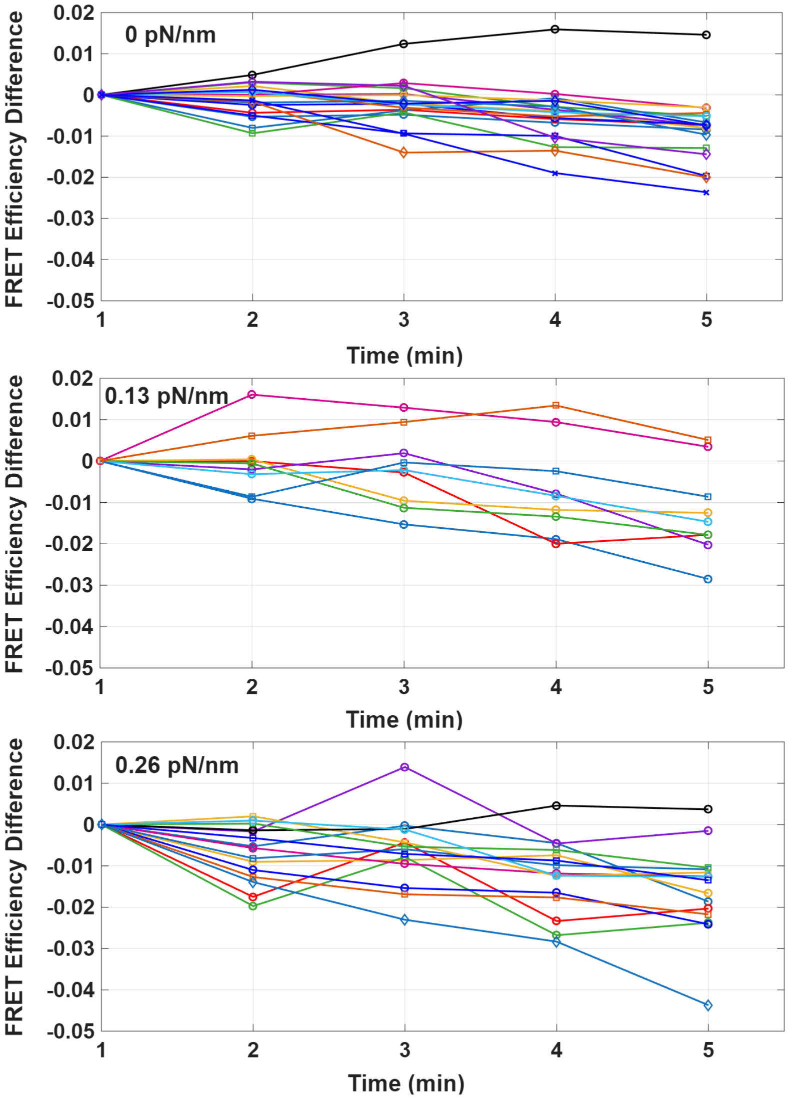

***Figure S8:*** *Plots of FRET efficiency difference as a function of time for each bead at the 3 trap conditions (No trap (0pN/nm), 0.13pN/nm and 0.26 pN/nm). Each trace represents a different bead.*

#### FRET efficiency difference after 5 min as a function of initial FRET efficiency

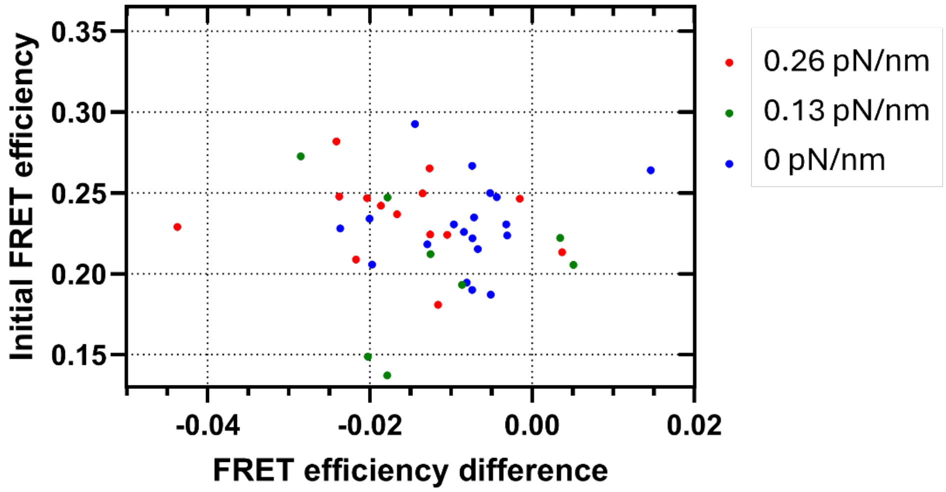

***Figure S9:*** *Initial FRET efficiency before turning on the laser trap vs efficiency difference between t=5 min and t=1min after applying the laser. Each dot corresponds to one bead, its color to the laser intensity applied: blue for no laser, green for medium trap stiffness, red for high trap stiffness.*

#### Vinculin intensity and FRET efficiency difference as a function of force for each bead

*Normalized AA intensity:*

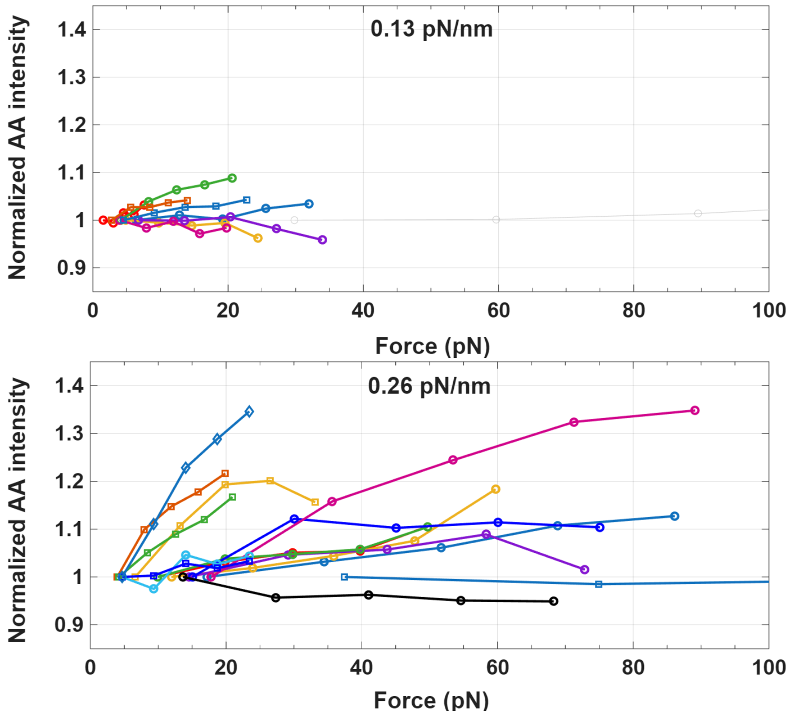

*FRET Efficiency Difference:*

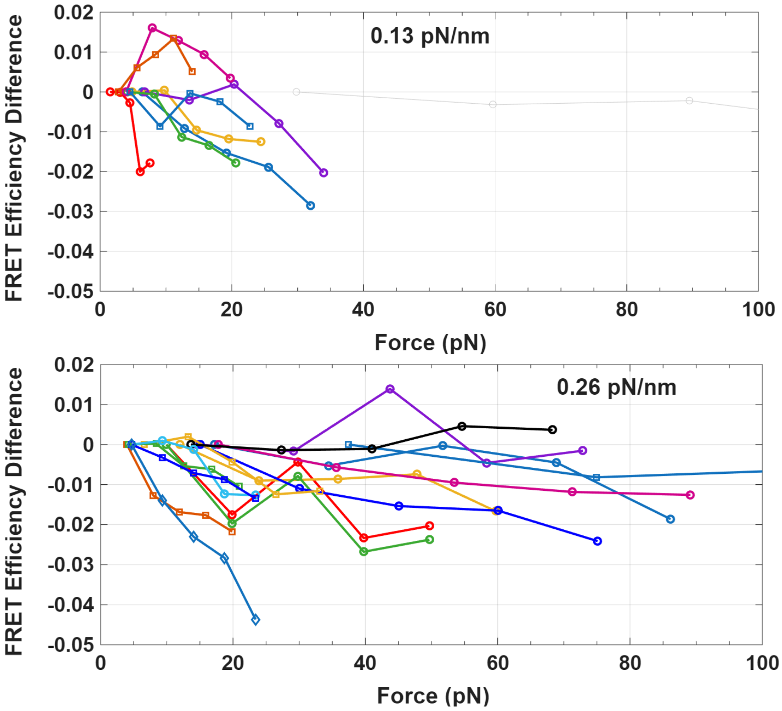

***Figure S10**: Normalized AA intensity and FRET efficiency difference as a function of force for each bead at 0.13 pN/nm and 0.26 pN/nm trap stiffness. Each trace represents a different bead starting at t= 1min. The first point corresponds to the force calculated based on the bead displacement between t= 0 and 1min. The points are connected in chronological order with force increasing as a function of time. Only data with displacement under 400nm (a maximum set by the linear range of the optical trap) are shown.*

#### Statistical Test Results for Figures 6a, 6b. 6c and 8a:

**Table S1:** Fig. 6a, Linear regression, Normalized AA intensity as a function of time.

| **Best-fit values** | **No trap** | **0.13 pN/nm** | **0.26 pN/nm** |
|---|---|---|---|
| Slope | -0.006617 | 0.006384 | 0.03443 |
| **Std. Error** | | | |
| Slope | 0.001269 | 0.001559 | 0.003176 |
| **Is the slope significantly non-zero?** | | | |
| P value | <0.0001 | 0.0002 | <0.0001 |

**Table S2:** Two-way ANOVA, Fig. 6a, Normalized AA intensity as a function of time, at different trap stiffnesses.

*Two-way ANOVA*

| **Factor** | **P value** |
|---|---|
| Time | 0.0013 |
| Trap stiffness | <0.0001 |
| Time*Trap stiffness | <0.0001 |

**Table S3:** Fig. 6a, Multiple comparisons, Normalized AA intensity at different trap stiffnesses.
*Tukey's multiple comparisons test*

| Timepoint | | P value |
|---|---|---|
| **2 min** | No trap vs. 0.13 pN/nm | 0.2959 |
| | No trap vs. 0.26 pN/nm | **0.0103** |
| | 0.13 pN/nm vs. 0.26 pN/nm | 0.0504 |
| **3 min** | No trap vs. 0.13 pN/nm | 0.0961 |
| | No trap vs. 0.26 pN/nm | **0.0020** |
| | 0.13 pN/nm vs. 0.26 pN/nm | **0.0159** |
| **4 min** | No trap vs. 0.13 pN/nm | **0.0450** |
| | No trap vs. 0.26 pN/nm | **0.0011** |
| | 0.13 pN/nm vs. 0.26 pN/nm | **0.0173** |
| **5 min** | No trap vs. 0.13 pN/nm | **0.0230** |
| | No trap vs. 0.26 pN/nm | **0.0003** |
| | 0.13 pN/nm vs. 0.26 pN/nm | **0.0206** |

**Table S4:** Fig. 6b, linear regression, FRET efficiency difference as a function of time.

| **Best-fit values** | **No trap** | **0.13 pN/nm** | **0.26 pN/nm** |
|---|---|---|---|
| Slope | -0.001805 | -0.002454 | -0.004025 |
| **Std. Error** | | | |
| Slope | 0.0002290 | 0.0005055 | 0.0003597 |
| **Is the slope significantly non-zero?** | | | |
| P value | <0.0001 | <0.0001 | <0.0001 |

**Table S5:** Fig. 6b, two-way ANOVA, FRET efficiency difference as a function of time, at different trap stiffnesses.
*Two-way ANOVA*

| Factor | P value |
|---|---|
| Time | <0.0001 |
| Trap stiffness | 0.0442 |
| Time*Trap stiffness | 0.0772 |

**Table S6:** Fig. 6b, multiple comparisons, FRET efficiency difference at different trap stiffnesses.
*Tukey's multiple comparisons test*

| Timepoint | | P value |
|---|---|---|
| **2 min** | No trap vs. 0.13 pN/nm | 0.9074 |
| | No trap vs. 0.26 pN/nm | **0.0193** |
| | 0.13 pN/nm vs. 0.26 pN/nm | 0.0938 |
| **3 min** | No trap vs. 0.13 pN/nm | 0.9939 |
| | No trap vs. 0.26 pN/nm | 0.2398 |
| | 0.13 pN/nm vs. 0.26 pN/nm | 0.4706 |
| **4 min** | No trap vs. 0.13 pN/nm | 0.9119 |
| | No trap vs. 0.26 pN/nm | **0.0360** |
| | 0.13 pN/nm vs. 0.26 pN/nm | 0.4360 |

| | | |
|---|---|---|
| **5 min** | No trap vs. 0.13 pN/nm | 0.5987 |
| | No trap vs. 0.26 pN/nm | 0.0854 |
| | 0.13 pN/nm vs. 0.26 pN/nm | 0.7344 |

**Table S7**: Fig. 6c, one-way ANOVA comparing the slopes of AA intensity and FRET efficiency at the 3 trap stiffnesses.

*One-way ANOVA followed by Tukey's multiple comparisons test*

| | | **P value** |
|---|---|---|
| **Normalized AA intensity** | One-way ANOVA | <0.0001 |
| **Slope of AA intensity multiple comparisons test** | No trap vs. 0.13 pN/nm | <0.0001 |
| | No trap vs. 0.26 pN/nm | <0.0001 |
| | 0.13 pN/nm vs. 0.26 pN/nm | <0.0001 |
| **FRET efficiency difference** | One-way ANOVA | <0.0001 |
| **Slope of efficiency difference multiple comparisons test** | No trap vs. 0.13 pN/nm | 0.0001 |
| | No trap vs. 0.26 pN/nm | <0.0001 |
| | 0.13 pN/nm vs. 0.26 pN/nm | <0.0001 |

**Table S8**: Fig. 8a, one-way ANOVA comparing the bead displacements at the 3 trap stiffnesses.

*One-way ANOVA followed by Tukey's multiple comparisons test*

| | **P value** |
|---|---|
| One-way ANOVA | 0.5817 |
| No trap vs. 0.13 pN/nm | 0.5634 |
| No trap vs. 0.26 pN/nm | 0.8350 |
| 0.13 pN/nm vs. 0.26 pN/nm | 0.8633 |
